# Robust associative learning is sufficient to explain structural and dynamical properties of local cortical circuits

**DOI:** 10.1101/320432

**Authors:** Danke Zhang, Chi Zhang, Armen Stepanyants

## Abstract

The ability of neural networks to associate successive states of network activity lies at the basis of many cognitive functions. Hence, we hypothesized that many ubiquitous structural and dynamical properties of local cortical networks result from associative learning. To test this hypothesis, we trained recurrent networks of excitatory and inhibitory neurons on memory sequences of varying lengths and compared network properties to those observed experimentally. We show that when the network is robustly loaded with near-maximum amount of associations it can support, it develops properties that are consistent with the observed probabilities of excitatory and inhibitory connections, shapes of connection weight distributions, overrepresentations of specific 3-neuron motifs, distributions of connection numbers in clusters of 3–8 neurons, sustained, irregular, and asynchronous firing activity, and balance of excitation and inhibition. What is more, memories loaded into the network can be retrieved even in the presence of noise comparable to the baseline variations in the postsynaptic potential. Confluence of these results suggests that many structural and dynamical properties of local cortical networks are simply a byproduct of associative learning.

## INTRODUCTION

With ever-increasing amounts of data on structure and dynamics of neural circuits, one fundamental question moves into focus: Is there an overarching principle that can account for the multitude of these seemingly unrelated experimental observations? For example, much is known about local connectivity in cortical circuits (see e.g. Supplementary Dataset). Some of the most salient connectivity features can be described based on the classes of excitatory and inhibitory neurons. At the level of pair-wise connectivity, it is known that the probabilities of excitatory connections are generally lower than those for inhibitory. Specifically, the majority of reported probabilities lies in the 0.10–0.19 range (interquartile range based on Supplementary Dataset) if the presynaptic cell is excitatory and in the 0.25–0.56 range for connections originating from inhibitory neurons. It is also known that the distributions of connection weights have stereotypic shapes with the majority of measured coefficients of variation (CV) in the 0.85–1.1 range for excitatory connections and slightly lower values for inhibitory, 0.78–0.96. At the level of connectivity within 3-neuron clusters, several ubiquitously overrepresented connectivity motifs have been discovered ^1–4^. Information becomes scarce as one considers larger clusters of neurons, but even here deviations from random connectivity have been reported for clusters of 3–8 neurons^2^. Similarly, many universal features characterize activity of neurons in local cortical networks. For example, individual neurons exhibit highly irregular spiking activity, resembling Poisson processes with close to one CV in inter-spike-intervals ^5–9^. Spike trains of nearby neurons are only marginally correlated, 0.04–0.15 ^10^, and, at the network level, spiking activity can be described as sustained, irregular, and asynchronous.

Two popular models of binary McCulloch and Pitts neuron networks ^11^ can individually explain some of the above experimental observations. The first model is based on the idea that to have a sustained and irregular activity, excitatory and inhibitory inputs to individual neurons in the network must be balanced ^12–17^. This model assumes that excitatory and inhibitory inputs are much larger than the threshold of firing, but their sum lies below the threshold, and firing is driven by fluctuations. The balanced model can produce realistic sustained and irregular spiking activity, however, by taking network connectivity as an input, it generally does not make predictions related to the network structure (but see ^18^). The second model, which we will refer to as the associative model, is based on the idea that synaptic connectivity is a product of associative learning ^19–23^. This model can explain many features of local cortical connectivity, but it does not necessarily produce sustained and irregular activity. We show that there is a biologically plausible regime, in which balanced and associative models converge. Therefore, with a single framework one can explain both structural and dynamical properties of cortical circuits and show that these properties emerge as a result of associative learning.

We pursue the idea that associative learning alone is sufficient to explain the above described properties of connectivity. With sensory information continuously impinging on the brain, neural circuits function in a state of perpetual change, recording some of the information in the form of long-term memories. In this process, individual neurons may be operating as independent learning units, constrained by functional and metabolic considerations, such as the requirement to store associative memories, tolerate noise during memory retrieval, and maintain low cost of the underlying connections ^24^, Figure 1A. In this study, we explore structural and dynamical properties of associative networks in the space of these constraints, and show that there is a unique region of parameters that can explain the above-described experimental observations. In this region, the network is loaded with close to maximum amount of associative memories which can be successfully retrieved even in the presence of significant amount of noise.

**Figure 1:**
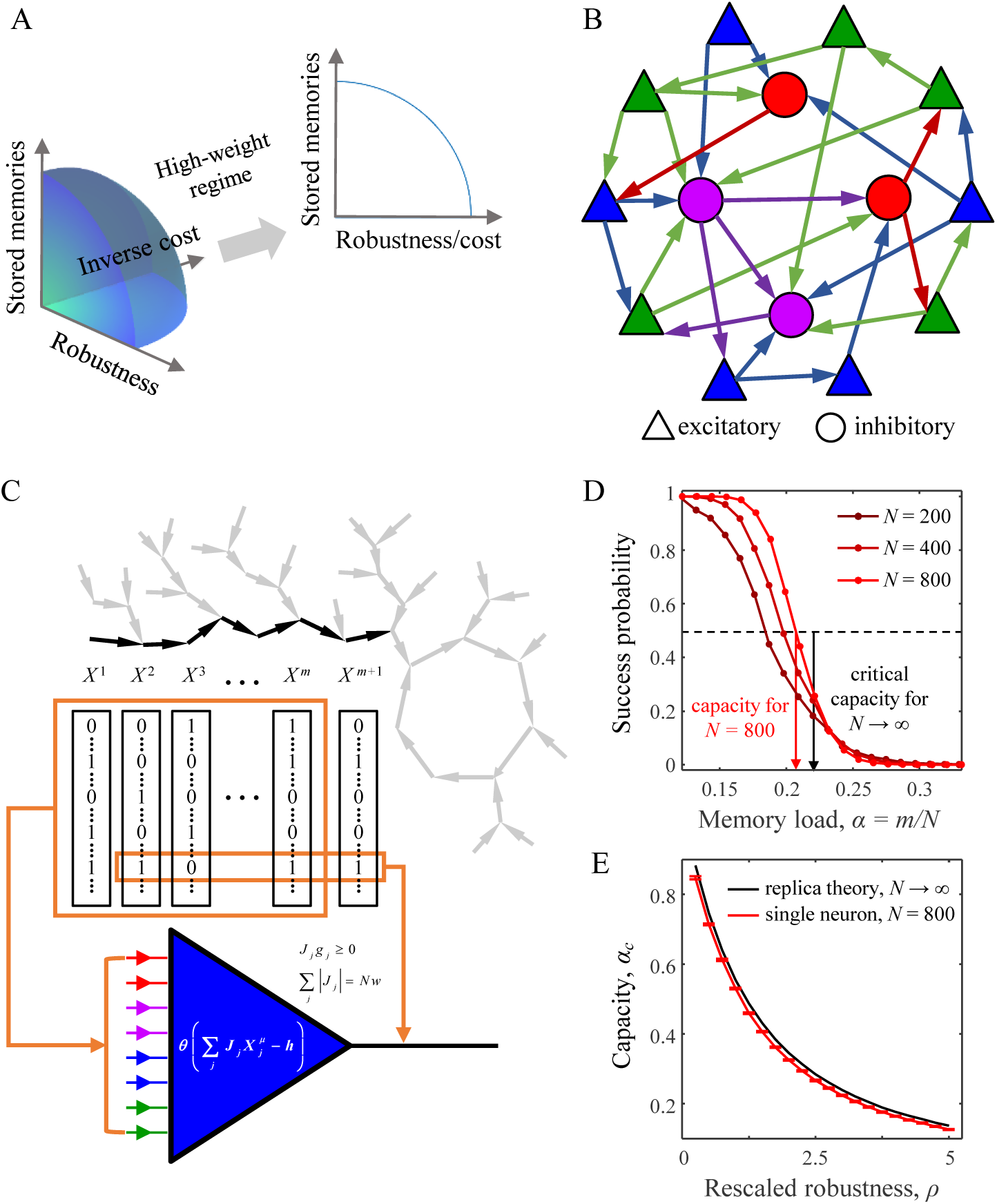
Associative memory storage in recurrent networks of excitatory and inhibitory neurons. **A.** Associative learning in the brain is expected to be constrained by functional and metabolic considerations, such as being able to store large amounts of memories, tolerate noise during memory retrieval, and have a low cost of the underlying connectivity. In the model, these three considerations are represented with memory load, *α*, robustness parameter, *κ*, and average absolute connection weight, *w*. We show that in the biologically plausible regime of high-weight, results of the model depend only on *α* and *κ/w*. **B.** Recurrent network of various classes (color) of all-to-all potentially connected excitatory and inhibitory neurons. Note that the arrows indicate actual (or functional) connections. **C.** Associative memory in the model is represented as a sequence (bold arrows), or an entire basin, of network states, *X*^1^ → *X*^2^ … → *X^μ^*. Vector *X^μ^* represents binary activities of individual neurons at time step *μ*. Each neuron in the network learns its corresponding set of input-output associations (orange boxes) by modifying the strengths of its input connections, *J_j_*, under the constraints on the signs and *l*_1_-norm of these connections. **D.** A neuron’s ability to learn an entire sequence of presented associations decreases with the sequence length, *m*. Memory storage capacity of the neuron, *α_c_*, (e.g. red arrow for *N* = 800) is defined as the fraction of associations, *m*/*N*, that can be learned with success probability of 50%. The transition from perfect learning to inability to learn the entire sequence sharpens with increasing *N* and approaches the result obtained with the replica theory in the limit of *N* → ∞ (black arrow). **E.** Capacity of a single neuron is a decreasing functions of the rescaled robustness, *ρ*. Error-bars indicate standard deviations calculated based on 100 networks.

## RESULTS

### Network model of associative learning

We use a McCulloch and Pitts neural network ^11^ to model a local cortical circuit in which *N_inh_* inhibitory neurons and (*N* – *N_inh_*) excitatory neurons are all-to-all potentially connected ^25,26^ (Figure 1B). Associative memories, in the form of temporal sequences of network states, are loaded in the network by modifying the weights of connections between neurons (see Online Methods and *SI* for details). In this process, individual neurons attempt to associate the inputs they receive from the network with predefined outputs (Figure 1C). Several biologically motivated constraints are imposed on the learning process ^24^. First, firing thresholds of neurons, *h*, do not change during learning. Second, the signs of input weights, *J*, that are determined by the excitatory or inhibitory identities of presynaptic neurons, do not change during learning (Dale’s principle) ^27^. Third, input connections of each neuron are homeostatically constrained to have a fixed average absolute weight, *w* ^28–30^. Fourth, each neuron must be able to retrieve the loaded associations even in the presence of noise in its postsynaptic potential. The maximum amount of noise a neuron has to tolerate is referred to as robustness parameter, *κ*.

Individual neurons in the model learn to associate the presented sequences of network states, and the probability of successfully learning the entire sequence decreases with the sequence length, *m*, or memory load *α* = *m/N* (Figure 1D). Memory load that can be successfully learned with the probability of 0.5 is termed the associative memory storage capacity of the neuron, *α_c_*. This capacity increases with the number of neurons in the network, *N*, and saturates in the *N →* ∞ limit at a value that can be determined with the replica theory ^20^ (see *SI*). Notably, this theoretical solution shows that in the high-weight regime (*Nwf* / *h* ≫ 1), the neuron’s capacity, as well as the shape of its connection weight distribution, depend on the combination of model parameters in the form of 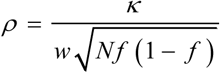, where *f* is the neuron’s firing rate (Figures 1A and S1). The meaning of this combination was elucidated by Brunel et al. ^21^, who pointed out that *ρ* can be viewed as a measure of reliability of stored associations to errors in the input. We would like add that *ρ* can also be viewed as a proxy for the ratio of robustness to fluctuations in postsynaptic potential. Following ^22^, we will refer to *ρ* as the rescaled robustness.

Motivated by this theoretical insight we set out to explore the possibility that local networks in the brain function in the high-weight regime. The average absolute connection weight, *w*, was previously estimated based on experimental data from various cortical systems ^24^, and the result shows that *Nwf* / *h* lies in the range of 4 - 38 (95% confidence interval) with the average of 14. A similar estimate based on the granule to Purkinje cell connectivity in rat cerebellum ^21^ also results in a relatively high value of this parameter, *Nwf* / *h* ≈ 150,000 × 0.1 mV × 0.0044/10 mV = 6.6. Therefore, high-weight regime may be a general attribute of local circuits, and we will show that this assumption is consistent with a large number of experimental measurements related to network structure and dynamics.

Figure 1E shows that the memory storage capacity of a single neuron is a decreasing function of rescaled robustness. This is expected, as increase in *ρ* can be thought of as an increase in the strength of the constraint on learning (robustness, *κ*), or as a decrease in the amount of available resources (absolute connection weight, *w*). Figure 1E also illustrates that at *N* = 800 single neuron capacity is already sufficiently close to its *N → ∞* limit, and therefore, network properties are not expected to change substantially if *N* is increased beyond 800 (Figure S2).

### Properties of neuron-to-neuron connectivity in associative networks

Next, we examined the properties of neuron-to-neuron connectivity in associative networks at different values of memory load and rescaled robustness. One of the most prominent features of connectivity is that substantial fractions of excitatory and inhibitory connections have zero weights, and therefore, connection probabilities are less than one (Figure 2A). The distributions of non-zero connection weights resemble the general shapes of unitary postsynaptic potential (uPSP) distributions, with a notable difference in the frequencies of strong connections. The former have Gaussian or exponential tails ^21,24^, while the tails of uPSP distributions are often much heavier ^1,31^. Several amendments to the associative model have been proposed to correct this discrepancy ^21,23^. Here, we would like to point out that heavy tails of experimental distributions can be reproduced within the associative model by considering networks of neurons with inhomogeneous properties, e.g. different values of *w* or *κ*.

**Figure 2:**
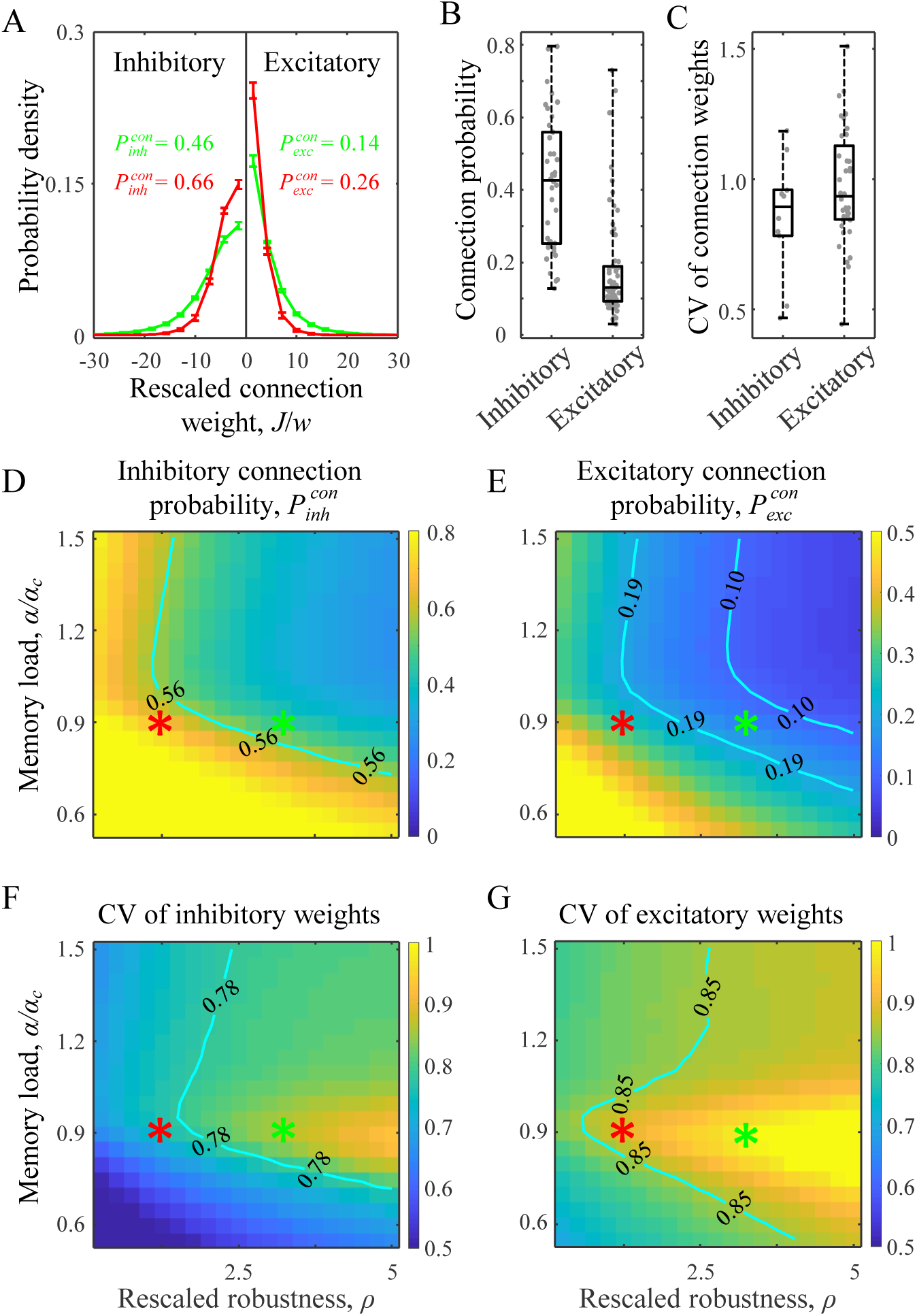
Properties of neuron-to-neuron connectivity in associative networks. **A.** Distributions of weights of inhibitory and excitatory connections for two parameter settings (red and green asterisks in D-G). Note that the distributions contain finite fractions of zero-weight connections. Error-bars indicate standard deviations (based on 100 networks). **B, C.** Connection probabilities and CVs of connection weights for inhibitory and excitatory connections reported in 87 studies describing 420 local cortical projections in mammals (Supplementary Dataset). Each dot represents the result of a single study averaged (with weights equal to the number of connections tested) over the number of reported projections. Maps of probabilities of inhibitory **(D)** and excitatory **(E)** connections as functions of rescaled robustness and relative memory load, i.e. load divided by the theoretical single neuron capacity at *N →* ∞. Inhibitory connection probability is higher than probability of excitatory connections in the entire region of considered parameters. **F, G.** Maps of coefficients of variation (CV) of non-zero inhibitory and excitatory connection weights as functions of rescaled robustness and relative memory load. Isocontour lines in the maps correspond to the interquartile ranges of experimentally observed connection probabilities and CVs shown in (B) and (C). Numerical results were generated based on networks of *N* = 800 neurons.

To compare connection probabilities and widths of non-zero connection weight distributions in associative networks with those reported experimentally, we compiled measurements of connection probabilities and CVs in uPSPs reported for local cortical circuits (Supplementary Dataset). Figures 2B, C show that the average inhibitory connection probability (based on 38 studies, 9,522 connections tested) is significantly higher (p < 10^−10^, two sample t-test) than the average probability for excitatory connections (67 studies, 63,020 connections tested), while CV of inhibitory uPSPs (10 studies, 503 connections recorded) is slightly lower than that for excitatory (36 studies, 3,956 connections recorded). Similar trends are observed in associative networks. Figures 2D, E show that connectivity in associative networks is sparse, with probabilities of excitatory non-zero connections lower than those for inhibitory connections in the entire considered range of memory load and rescaled robustness. Probabilities of both connection types are decreasing with increasing *ρ*. This is expected because an increase in *ρ* can be achieved by lowering *w*, which is equivalent to reducing the amount of resources needed to make connections. Isocontours in Figures 2D, E demarcate the interquartile ranges of connection probability measurements shown in Figure 2B. There is a region in the *α–ρ* space of parameters in which both excitatory and inhibitory connection probabilities are in general agreement with the experimental data. Also, consistent with the experimental measurements, CVs of excitatory weights in associative networks are slightly larger than those for inhibitory weights (Figures 2F, G), and there is a wide region in the *α–ρ* space of parameters in which these values match the experimental data shown in Figure 2C.

### Higher order structural properties of associative networks

In addition to specific properties of neuron-to-neuron connectivity, local cortical circuits are known to have non-random patterns of connections in subnetworks of three and more neurons ^1–4^. To determine whether associative networks can reproduce some of the known features of higher-order connectivity, we first examined the statistics of connectivity motifs within subnetworks of three excitatory neurons. There are 13 distinct types of connected 3-neuron motifs (Figure 3A). Under/over-expressions of these motif types were quantified with the normalized *z-*scores, which range from −1 to 1 and are negative/positive for motifs that appear less/more frequently than what is expected by chance (see Methods). Profiles of normalized *z-*scores in associative networks were compared with data from the Blue Brain project ^4^. To gauge the extent of similarity of these profiles we calculated Root-Mean-Square (RMS) difference in the normalized *z-*scores for various values of load and rescaled robustness (Figure 3B). The results show that there is a region of parameters in which associative networks produce profiles similar to the Blue Brain project data. In particular, motifs 3, 4, 10, and 11 appear to be ubiquitously overexpressed in the region of parameters outlined by the isocontour in Figure 3B, which is in agreement with the Blue Brain project data (see e.g. green line in Figure 3A) and data from other experiments ^1,2,4^.

**Figure 3:**
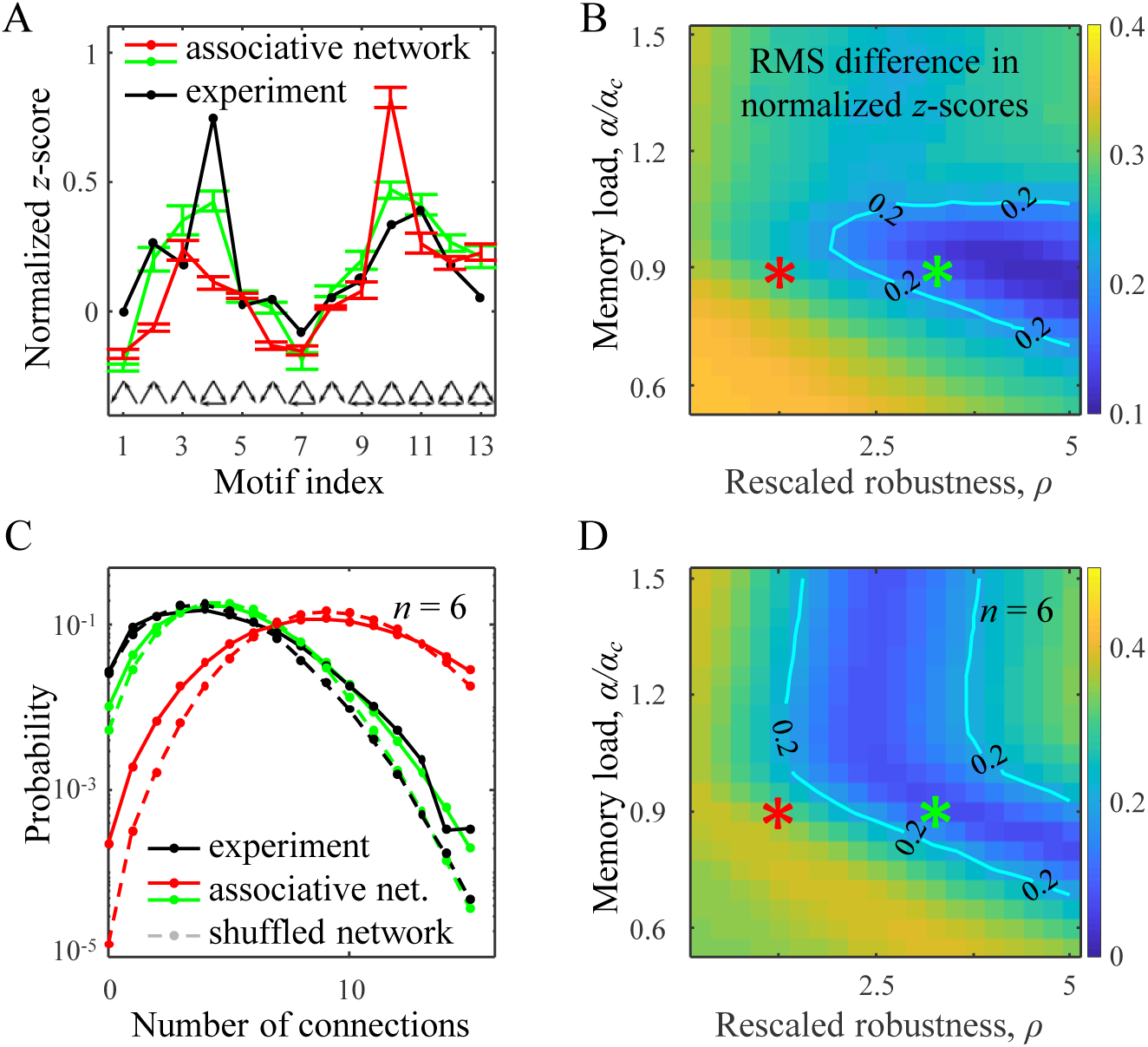
Higher-order structural motifs in associative networks. **A.** Normalized *z-*score of 13 connected 3-neuron motifs in excitatory subnetworks indicate over-and under-expressions of these structures in comparison to the chance level (see Methods for details). Red and green curves show results for the parameter settings specified by the red and green asterisks in (B). Black curve is the result from the Blue Brain project ^4^. Error-bars indicate standard deviations (100 networks). **B.** Root-mean-square (RMS) difference between normalized *z-*scores obtained in associative networks and in the Blue Brain project. Isocontour line, demarcating a region of reasonably good solutions, is drawn as a guide to the eye. **C.** Distributions of non-zero connection numbers in clusters of 6 excitatory neurons in associative networks. Solid red and green lines illustrate distributions obtained in associative networks for the parameter settings indicated by the red and green asterisks. Solid black curves indicate the corresponding results for local cortical networks based on electrophysiological measurements ^2^. Dashed lines show distributions in randomly shuffled networks (see Methods for details). **D.** Maps of *l*_2_ distances between connection number distributions in associative and cortical networks ^2^. Similar results for clusters of 3–8 neurons are shown in Figure S5. Numerical results of the associative model were generated based on networks of *N* = 800 neurons.

Deviations from random connectivity have also been detected in subnetworks of 3 to 8 excitatory neurons by comparing distributions of observed connection numbers with those based on randomly shuffled connectivity ^2^. This comparison revealed that the experimental distributions have heavier tails, which is indicative of clustered connectivity (black lines in Figures 3C and S5). This trend was first reproduced by Brunel ^22^ who considered an associative network of excitatory neurons at capacity. Our model shows that there is a single region of parameters *α* and *ρ* in which qualitative agreement is obtained simultaneously for all subnetworks from 3 to 8 neurons (Figures 3D and S5).

### Dynamical properties of associative networks

Dynamics in associative networks depends strongly on the values of memory load and rescaled robustness. At small values of *ρ*, network dynamics quickly terminates at a fixed point in which all neurons are silent (Figure 4A1, red). When *ρ* is high, associative networks can have long-lasting intrinsic activity, often ending up in a limit cycle of non-zero length. To quantify this behavior, we measured the average number of steps taken by the network to reach a limit cycle or a fixed point from a random initial state (Figure 4A2). The results show that the duration of transient dynamics increases exponentially with memory load and rescaled robustness. Even for moderate values of these parameters, the average length of transient activity can be of the order of network size, *N* (contour in Figure 4A2).

**Figure 4:**
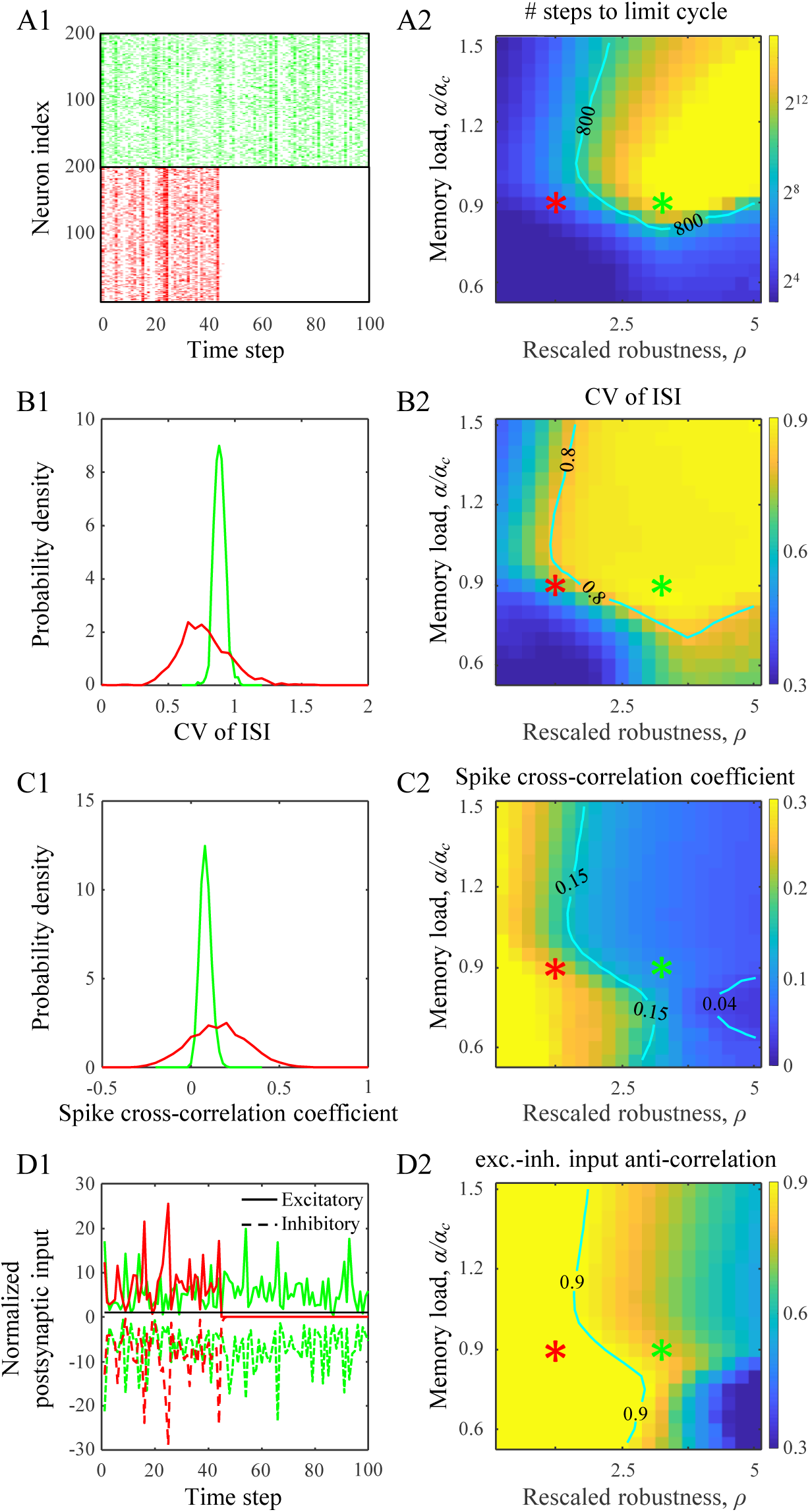
Dynamical properties of associative networks. **A1**. Two examples of spike rasters for associative networks parametrized as indicated with red and green asterisks in (A2). Dynamics at low values of rescaled robustness (red) quickly terminates at a quiescent state. **A2**. Map of the duration of transient dynamics as a function of rescaled robustness and relative memory load (see Methods for details). Note that at high levels of rescaled robustness and memory load, associative networks have long-lasting, transient activity. Isocontour line is drawn as a guide to the eye. **B1**. Distributions of CV in inter-spike-intervals (ISI) for the two parameter settings. Note that the average CV value increases with *ρ*. **B2.** Map of the average CV of ISI as a function of rescaled robustness and relative memory load. Isocontour line demarcates a region of high CV values that are in general agreement with experimental measurements. **C.** Same for cross-correlation coefficients of neuron spike trains. **D.** Same for anti-correlation coefficient of excitatory and inhibitory postsynaptic inputs received by a neuron. Note that the inputs are normalized by the firing threshold. For the selected parameter configurations, excitatory and inhibitory inputs are tightly balanced (large negative anti-correlation) despite large fluctuations. Maps in (A2, B2, C2, and D2) were generated based on networks of *N* = 800 neurons by averaging the results over 100 networks and 100 trials for each network and parameter setting.

Individual neurons in associative networks are capable of producing irregular spiking activity, degree of which can be quantified with CV of inter-spike-intervals (ISI) (Figure 4B). According to this measure, neurons exhibit greater irregular activity when memory load and rescaled robustness are high, with CV of ISI values saturating at around 0.9. This is consistent with the range of CV of ISI values reported for different cortical systems, 0.7–1.1 ^5–9^. To examine the extent of synchrony in neuron activity we calculated spike train cross-correlation coefficients for pairs of neurons (Figure 4C). The results show that increase in *ρ* leads to a more asynchronous activity, which can be explained by the reduction in connection probability (Figures 2D, E) and, consequently, reduction in the amount of common input to the neurons. For *ρ* > 2.5, the values of cross-correlation are consistent with experimental data 0.04–0.15 ^10^ (interquartile range derived from 26 studies).

Irregular, asynchronous activity can result from balance of excitation and inhibition ^12,13^. In the balanced state, the magnitudes of excitatory and inhibitory potentials are typically much greater than the threshold of firing, and, due to a high degree of correlation in these potentials, firing is driven by fluctuations. Consistent with this, the average excitatory and inhibitory inputs in the associative model are much greater than the firing threshold and are tightly anti-correlated (Figures 4D and S7). The degree of anti-correlation decreases with rescaled robustness as the network connectivity becomes sparser. Experimentally, it is difficult to measure anti-correlations of excitatory and inhibitory inputs within a given cell, but such measurements have been performed in nearby cells. The resulting anti-correlations, ~0.4 ^32,33^, are somewhat below the values observed in associative networks. However, this is expected, as between-cell anti-correlations are likely to be weaker than within-cell anti-correlations.

### Cortical circuits are loaded with associative memories close to capacity and can tolerate noise comparable to the baseline variations in postsynaptic input during memory retrieval

Parameter regions described in Figures 2–4, lead to structural and dynamical properties consistent with the experimental observations and have a non-empty intersection. In this biologically plausible region of parameters, associative networks behave qualitatively similar to local cortical circuits. Figure 5A shows the intersection of parameter regions (green dashed line) for the excitatory and inhibitory connection probabilities (red), 3-neuron motifs (green), connections in 3–8 neuron clusters (blue), and duration of transient activity (cyan). The remaining features, i.e. CV of connection weights, CV of ISI, spike cross-correlation coefficient, and excitatory-inhibitory balance, are not shown in Figure 5A both to avoid clutter and because they do not impose additional restrictions on the intersection region. In the biologically plausible region of parameters, individual neurons are loaded with relatively long sequences of network states (0.2*N* for the green asterisk in Figure 5A), yet it is not clear if the associations learned by individual neurons assemble into memory sequences that can be successfully retrieved at the network level.

**Figure 5:**
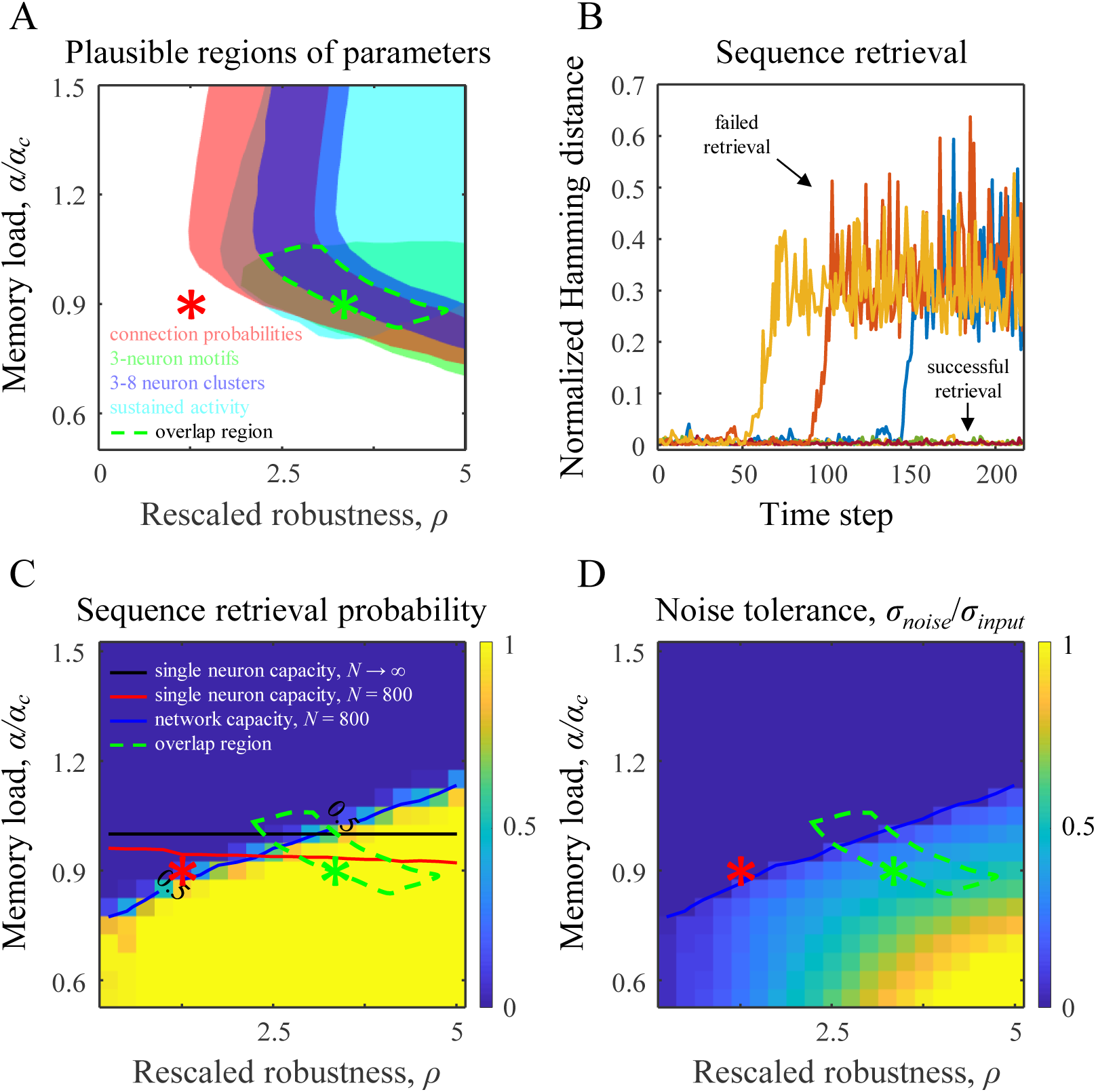
Values of rescaled robustness and memory load identified based on structural and dynamical properties of local cortical networks are consistent with the functional requirement of robust retrieval of stored memories. **A.** Region of parameters (dashed green line) that leads to a general agreement with the experimentally observed excitatory and inhibitory connection probabilities (red), excitatory 3-neuron motifs (green), 3–8 excitatory neuron clusters (blue), and sustained, irregular, asynchronous spiking activity (cyan). **B.** Retrieval of memory sequences in the absence of noise. The network is loaded with a memory sequence that it attempts to learn. Retrieval process is initialized at the start of the sequence, and deviations of subsequent network states from the loaded states are quantified with the Hamming distance normalized by the network size, *N*. The sequence is said to be successfully retrieved if the deviations are small. **C.** Success probability of sequence retrieval as a function of rescaled robustness and relative memory load. Blue line, corresponding to the success probability of 0.5, defines the network capacity. Single neuron critical capacity for *N* → ∞ (black line) and capacity for *N* = 800 (red line) are shown for reference. Dashed green contour is the overlap region from (A). **D.** Noise level, *σ_nοise_*, that can be tolerated by the associative network during memory retrieval. Map shows the relative noise level, σ*_noise_*/σ*_input_*, corresponding to the retrieval probability of 0.5. Note that in the identified parameter region, the network can tolerate noise that is comparable to the standard deviation in postsynaptic input, σ*_input_*.

To examine this question we first tested memory retrieval in the absence of noise. For this, we initialized the network state at the beginning of the loaded sequence and monitored playout of the memory. The sequence is said to be retrieved successfully if the network states during the retrieval do not deviate substantially from the loaded states (Figure 5B). In practice, there is no need to precisely define the threshold amount of deviation. This is because for large networks, e.g. *N* = 800, the Hamming distance between the loaded and retrieved sequences either remains within ~ 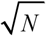 or diverges to ~N. Figure 5C shows the probability of successful memory retrieval as function of memory load and rescaled robustness. The transition from successful memory retrieval to inability to retrieve the entire loaded sequence is relatively sharp, making it possible to define network capacity, analogously to single neuron capacity, as the sequence length for which the success rate in memory retrieval equals 0.5 (blue line in Figure 5C). Network capacity deviates from single neuron capacity, but this difference is expected to decrease with network size. Interestingly, the biologically plausible region of parameters is centered on the single neuron capacity curve, implying that individual cortical neurons are loaded with associations close to their capacity. What is more, the biologically plausible region of parameters lies almost entirely below the network capacity curve, indicating that loaded memory sequences can be retrieved with high probability in the absence of noise.

To assess the degree of robustness of the memory retrieval process, we monitored memory playout in the presence of postsynaptic noise. In this experiment, random Gaussian noise of zero mean and standard deviation *σ_noise_* was added independently to all neurons at every step of the retrieval process. Network tolerance to noise was defined as *σ_noise_* value that results in the memory retrieval probability of 0.5. Figure 5D shows the map of noise tolerance normalized by the baseline variations in postsynaptic input during memory retrieval, *σ_noise_/σ_input_* (see *SI*). The latter represents standard deviation in postsynaptic input in the absence of noise. We note that the biologically plausible region identified on the basis of structural and dynamical properties of cortical networks (green contour in Figure 5D) has a non-zero overlap with the area in which memory retrieval is robust to noise. In this domain the network can tolerate high noise-to-input ratios (up to 0.5), which serves as an independent validation of the associative model in terms of the hypothesized network function.

## DISCUSSION

Our results suggest that local circuits of the mammalian brain operate in a high-weight regime in which individual neurons are loaded with associative memories close to their capacity (Figure 5C) and the network can tolerate relatively large amounts of postsynaptic noise during memory retrieval. In this regime, many structural and dynamical properties of associative networks are in general agreement with experimental measurements from various species and brain regions. It is important to point out that, due to large uncertainties in the reported measurements, we did not attempt to quantitatively fit the associative model to the data. The uncertainties originate from natural variability of network features across individuals, brain areas, and species, and are confounded by experimental biases and measurement errors. Instead, we rely on a large body of qualitative evidence to support our conclusions. These evidence include (1) sparse connectivity, with probability of excitatory connections being lower than that for inhibitory connections, (2) distributions of non-zero connection weights with CVs of excitatory and inhibitory weights being close to 1, (3) overrepresentations of specific 3-neuron motifs, (4) distributions of connection numbers in subnetworks of 3–8 neurons showing clustering behavior, (5) sustained, irregular, and asynchronous firing activity with close to 1 CV of ISI and small positive cross-correlation in neuron activity, and (6) balance of excitatory and inhibitory postsynaptic potentials. Many of these features have been separately reported in various formulations of the associative model ^20–24^. Here, we show that with a single set of model parameters it is possible to account for these features collectively. What is more, the identified set of model parameters overlaps with the region in which loaded memories can be successfully recalled even in the presence of postsynaptic noise (Figure 5D), providing an independent functional validation of the theory.

Several discrepancies between the results of the associative model and experiment are worth mentioning. First, the model does not produce overexpression of bidirectional connections observed in some experiments ^1,34,35^. This can be amended by including point attractors ^22^ in addition to temporal sequences of network states considered in this study. However, since not all experiments report overexpression of bidirectional connections, this feature may be area specific and/or dependent on the distance between neurons ^4^. Second, associative networks did not produce a good agreement with the distribution of inhibitory 3-neuron motifs reported by the Blue Brain project ^4^. We believe that this discrepancy can be attributed to a large diversity of inhibitory neuron population, which is not captured by the presented homogeneous model. Third, our theory does not produce long-tailed distributions of connection weights observed in many experiments ^1,31^. Several ways to amend this discrepancy have been previously discussed ^21,23^. It is also clear that introducing inhomogeneity in neuron parameters can broaden the tail of the connection weight distribution.

Because local cortical circuits function in the high-weight regime, *Nwf* ≫ *h*, the average excitatory and inhibitory postsynaptic inputs are significantly greater than the threshold of firing (Figure S7). In the identified region of memory load and rescaled robustness, e.g. for the green asterisk in Figure 5, these potentials in magnitude exceed the threshold of firing by factors of 6.3 and 7.8 respectively (Table S1). In this regime, excitatory and inhibitory potentials are strongly anti-correlated (Figure 4D2), which is reminiscent of the balanced state described by many authors ^16,32,33,36–39^. We note, however, that there is a difference in how balance of excitatory and inhibitory potentials is realized in the associative vs. balanced networks. The difference originates from scaling of synaptic weight with network size. In associative networks, synaptic weight is inversely proportional to *N*, while in balanced networks inverse proportionality to 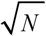 is assumed. In the former model, the average excitatory and inhibitory postsynaptic inputs to a neuron remain unchanged as the network size increases, and balance is the consequence of the high-weight regime, while in the latter model balance emerges with increasing *N* as postsynaptic potentials diverge, which is unsettling. On the other hand, Rubin et al. ^18^ argue that due to the above scaling difference, synaptic connections in the associative model are weaker, and the network is unstable to large, *O*(*h*), noise arising from processes within neurons (e.g. threshold fluctuations). We agree that susceptibility of associative networks to this type of noise is a concern for infinitely large systems. However, there is no biological data on scaling of noise with network size, and having *O*(*h*) noise may be unrealistic. More importantly, since local brain networks are finite, robustness to this type of noise can always be achieved by increasing *w*, i.e. in the high-weight regime. For example, an associative network of *N* = 800 neurons, configured at the green asterisk in Figure 5, can tolerate CVs in threshold fluctuations of up 1.1 (see Figure S7). Aside from the issue of robustness to *O*(*h*) noise, we show that in the high-weight regime results of the balanced and associative models become independent of the details of scaling, and converge to the same solution (see *SI* text and Figure S1). Therefore, associative learning in both models will lead to networks with identical structural and dynamical properties.

## METHODS

### Associative model

We consider an all-to-all potentially connected neural network of *N*_inh_ inhibitory and (*N* – *N*_inh_) excitatory McCulloch and Pitts neurons ^11^ involved in an associative learning task, Figure 1B. Biological motivations and assumptions associated with this model have been previously described ^24^. Here we only give a concise description of the model, as a more detailed account is provided in *SI*. Neurons in the model may belong to various classes, defined by their excitatory or inhibitory nature, characteristic firing probabilities, homeostatic constraints, and robustness to noise (defined below). The state of the network at time step *μ* is described by a vector of binary (0 or 1) activities of all neurons, *Χ^μ^*. The network is loaded with a predefined temporal sequence (or a basin) of *m+1* network states, *X*^1^ → *X*^2^ *→* … *X*^m+1^, and individual neurons, independently from one another, attempt to robustly learn to associate inputs and outputs derived from this sequence, Figure 1C. We assume that the network states to be learned are uncorrelated across neurons and time. Learning in the network is mediated by changing neuron connection weights, {*J_ij_*} (connection from neuron *j* to neuron *i*), in the presence of several biologically inspired constraints. (1) Input connection weights of each neuron are sign-constrained to be non-negative if the presynaptic neuron is excitatory and non-positive if it is inhibitory. (2) Input weights of each neuron are homeostatically constrained to have a predefined *l*_1_-norm. (3) Each neuron must attempt to learn its associations robustly, so that they can be recalled correctly in the presence of a given level of postsynaptic noise. The model can be summarized as follows (see Figure 1C):

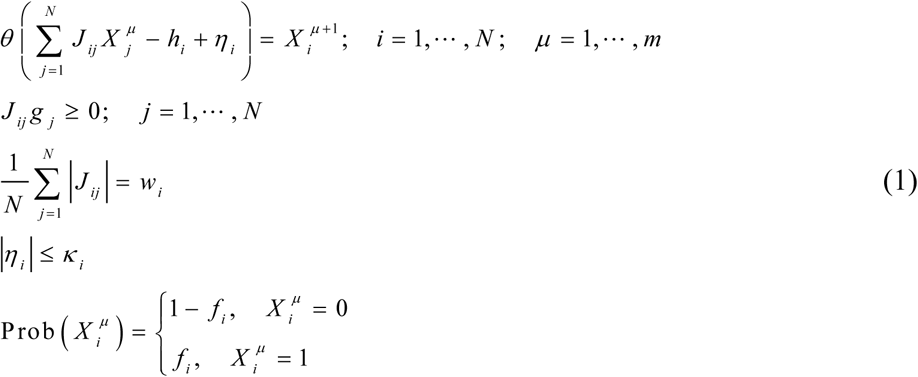

In these expressions, *θ* denotes the Heaviside step-function, *h_i_* is firing threshold, and *η_i_* denotes postsynaptic noise which is bounded by the robustness parameter *κ_i_*, i.e. |*η_i_*| < *κ_i_*. To enforce sign-constraints on connection weights we introduce parameter *g_j_*, which equals 1 if the presynaptic neuron *j* is excitatory and −1 if it is inhibitory. Parameter *w_i_*, referred to as the average absolute connection weight, is introduced to impose the *l*_1_-norm constraint on the neuron’s input connection weights. Binary neuron states, 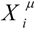, are randomly drawn from neuron-class dependent Bernoulli probability distributions: 0 with probability 1 *– f_i_* and 1 with probability *f_i_*.

Given a set of associations, the problem outlined in Eqs. (1) may be feasible and have multiple solutions *{J_ij_}*, or non-feasible, in which case no network can satisfy all the constraints of the problem. In the first case, the solutions region is nonempty, and we must employ additional considerations to limit the results to a single, unique solution. We did this by choosing the solution that minimizes the *l*_2_-norm of input connection weights of every neuron. In the non-feasible case, similar to what is done in the formulation of the Support Vector Machine problem ^40^, for every neuron, we minimized the sum of deviations between the not robustly learned associations and their corresponding margin boundaries (see *SI* for details).

The above model is governed by the following parameters (Table S1): number of neurons in the network, *N*, fraction of inhibitory neurons, *N_inh_/N*, firing probabilities of neurons in the associative sequence {*f_i_*}, robustness parameters of neurons {*κ_i_*}, their average absolute connection weights, {*w_i_*}, and the memory load, *α* = *m/N*. All numerical simulations presented in the main text were performed a homogeneous associative model, in which all excitatory and inhibitory neurons have the same firing threshold, *h*, average firing rate, *f*, constraints, *w* and *κ*, and memory load, *α*. We set the fraction of inhibitory neurons to *N_inh_/N* = 0.2, the firing rate to *f* = 0.2, and *Nwf* / *h* = 14, which is similar to what was described previously ^24^. We confirmed that the results are not sensitive to small changes in these parameters. The network size was chosen to be *N* = 800. In *SI* we reproduce the results for networks of *N* = 200, *N* = 400, and *N* → ∞, to illustrate that our conclusions do not depend on the exact value of *N* (Figures S2-4 and S6).

### Numerical and theoretical solutions of the associative model

Because individual neurons in the network learn independently of one another, the problem of sequence learning by a network can be solved individually for each neuron. In addition, since the model outlined in Eqs. (1) is convex, solution for each neuron can be obtained numerically with the methods of convex optimization ^41^ (see *SI* for details). Briefly, sequences of random, binary, and independent network states were generated with firing probability *f*. Individual neurons in the network were trained separately on their corresponding input-output associations extracted from these sequences. Probabilities of successful learning for single neurons was calculated by presenting the network with associative sequences of varying lengths (100 times for each length). Memory capacity of a single neuron is defined as the length of sequences the neuron can learn with success probability of 0.5 (Figure 1D).

In the limiting case of large networks, *N* → ∞, success probability abruptly changes from one to zero with increasing memory load, and neuron’s associative capacity is referred to as critical. In this limit, results of the model at critical capacity were obtained with the replica theory ^42,43^ (see *SI* for details).

Network capacity is defined based on the probability of successfully retrieving loaded associative sequences in the absence of noise (Figure 5C).

### Numerical simulations of structural and dynamical properties of associative networks

To examine properties of associative networks for different values of memory load and rescaled robustness, we trained 100 networks for every pair of these parameters, and used the resulting connection weight matrices to characterize network structure and dynamics.

Probability densities of connection weights (Figure 2A) and CV of connection weights (Figures 2F, G) were calculated after excluding small weights, i.e. weights with magnitudes less than 5*h/N*. We visually confirmed that this threshold encompasses the central peak in the connection weight distribution, associated with small excitatory and inhibitory weights. We also confirmed that the network properties described in the main text are not sensitive to the exact value of this parameter in the *5h/N –* 20*h/N* range. To analyze structural properties of associative networks (Figures 2D, E, and 3), weight matrices were converted into adjacency matrices by setting the small weights to zero and the remaining weights to 1. Numbers of 3-neuron motifs (Figures 3A,B) were calculated with the Brain Connectivity Toolbox ^44^. Frequencies of 13 connected 3-neuron motifs, *n_i_*, in subnetworks of excitatory neurons were compared with corresponding frequencies in subnetworks with randomly shuffled connections, 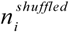. We used normalized *z-*scores, 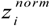, as defined in ^4^, to characterize the degrees of over- and under-expression of motif types:

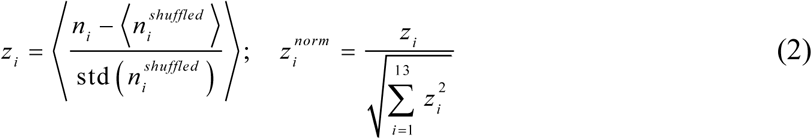

Here, outer angle brackets in the first equation denote averaging over 100 associative networks, angle brackets in the numerator represent averaging over a set of 50 randomly shuffled versions of a given associative network, and std denotes standard deviation over the set of shuffled networks. Normalized *z-*scores lie in the range of −1 to 1, and a positive *z-*score indicates that the observed number of motifs of a given type is larger than that expected by chance.

Dynamical properties of associative networks shown in Figure 4 were averaged over 100 networks and 100 random initial states for each network.

### Data and code availability

The dataset of connection probabilities and strengths used in this study was compiled from articles published in peer-reviewed journals from year 1990 to 2016. Specifically, we targeted publications reporting connection probabilities between neurons and uPSP amplitudes in local circuits of the mammalian brain. Using Google Scholar queries based on the above criteria, we initially identified 152 articles describing a total of 856 projections. Later we limited our analyses to experiments in which recordings were made in the neocortex, from at least 10 pairs of neurons located in the same layer and separated laterally by less than 100 μm. We also limited the analyses to normal, juvenile or adult animals (no younger than P14 for mouse and rat, and older than that for ferret, cat, monkey, and human). After imposing these limits, the numbers of publications and projections reduced to 87 and 420 respectively. These projections are included in the Supplementary Dataset. Inhibitory and excitatory connection probabilities and CVs based on these projections are shown in Figures 2B, C, in which to reduce bias we averaged the results of individual studies reporting multiple projections of a given type.

MATLAB code for generating theoretical (replica) and numerical (convex optimization) solutions of the associative model is available at https://github.com/neurogeometry/AssociativeLearning.

## ACKNOWLEDGEMENTS

This work is supported by the AFOSR grant FA9550-15-1-0398 and the NSF grant IIS-1526642.

## SUPPLEMENTARY INFORMATION

This Supplementary Information describes the model of associative memory storage by a recurrent network considered in the main text. The model incorporates a number of constraints motivated by the experimental data on connectivity in the cerebral cortex. The model is solved theoretically with the replica method ^1,2^ in the limit of infinite network size, and numerically with methods of convex optimization ^3^ for large, but finite networks. The model gives rise to a comprehensive list of predictions regarding the structure and dynamics of neural networks. These predictions are consistent with a large number of experimental studies of connectivity in local cortical circuits. Other models, including only some of the constraints considered in this study, have been previously described ^4–9^.

### General assumptions and approximations

This model is based on a number of assumptions and approximations, some of which have been previously discussed ^7,8^:

- We consider a recurrent network of *N* all-to-all potentially connected ^10,11^ McCulloch and Pitts neurons ^12^. Each neuron may belong to one of several classes defined by the values of the firing threshold, *h*, firing probability, *f*, average absolute weight of input connections, *w*, and robustness parameter, *κ*.
- The network is presented with associative memory sequences consisting of synchronous network states that are independent across neurons and time.
- Individual neurons in the network attempt to learn, independently from one another, input-output associations derived from these sequences.
- Each neuron learns by modifying its input connection weights, *J*, in the presence of constraints on the signs and *l*_1_-norm of these connections. Parameters *h*, *f, w*, and *κ* remain fixed during learning.
- In numerical simulations, when the memory load is subcritical and a neuron is presented with a feasible learning problem, we choose the solution with the minimum *l*_2_-norm of input connection weights. For a higher memory load, when the associative learning problem is non-feasible, we choose the solution that minimizes the total error associated with the erroneously learned associations.
- For the replica theory calculations performed in the limit of *N → ∞*, we consider two specific scenarios of scaling of model parameters with *N*. We show that in the biologically plausible limit of parameters, both scenarios lead to the same results. We note, that scaling assumptions are not required for numerical simulations performed for finite *N*.
- Memory retrieval is deemed to be successful if the retrieved sequence does not deviate significantly from the learned associative sequence.

### Learning capacity of a single neuron with a fixed threshold, binary (0, 1) input/output, robustness to noise, sign constraints, and multiple equality constraints

We consider a single neuron receiving *N* potential inputs from the network (enumerated with index *j*). The neuron attempts to learn a set of *m* binary (0, 1) input-output associations, 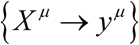, in the presence of postsynaptic noise, *η*, and constraints on its input connection weights, *J_j_*:

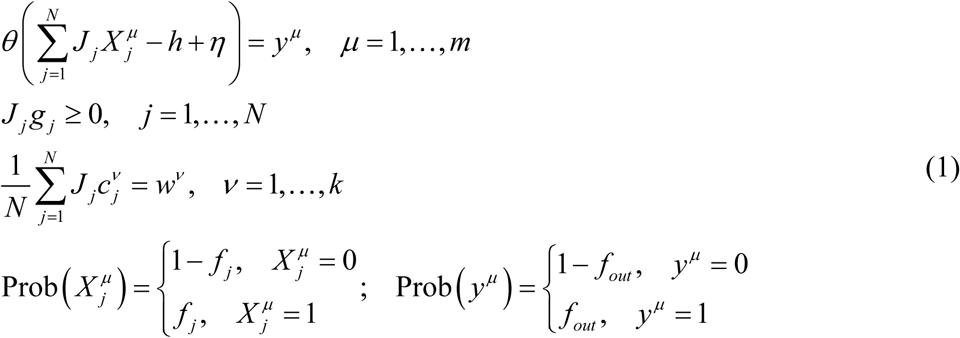

In these expressions, *θ* denotes the Heaviside step-function and *h* is the firing threshold. Inputs 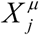 and outputs *y^μ^* are randomly and independently drawn from distinct Bernoulli distributions, in which the probabilities of having 1 are denoted with *f_j_* and *f_out_* respectively. To enforce sign-constraints on connection weights we introduced parameters *g_j_*, which equal 1 for excitatory and −1 for inhibitory inputs. Parameters 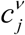 and *w^v^* define *k* linear equality constraints.

We assume that the associations must be learned robustly so that they can be successfully recalled in the presence of postsynaptic noise, *η*, |*η*| ≤ *κ*. Parameter *κ ≥* 0 is referred to as the robustness parameter. After eliminating *η* from Eqs. (1), the problem can be rewritten as:

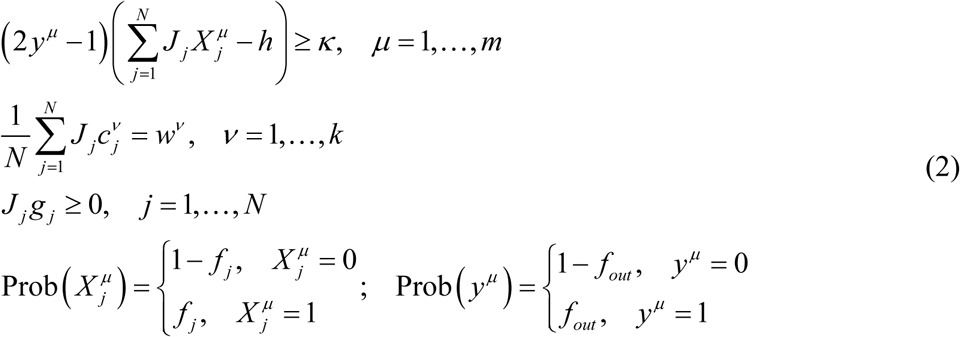

### Additional assumptions required for the replica theory calculation

We assume that *N* is large, while *m/N*, *f_j_* and *f_out_* and 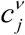. are *O*(1) (of order 1 with respect to *N*). Total input to a neuron can be expressed in terms of the input averaged over the associations, *X*, plus a deviation from the average:

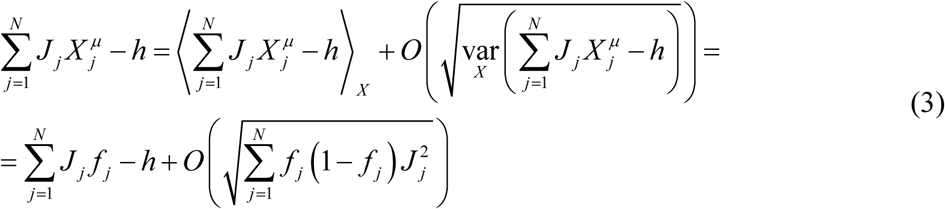

With this, the associative learning problem of Eqs. (2) can be separated into two categories based on the value of *y^μ^*:

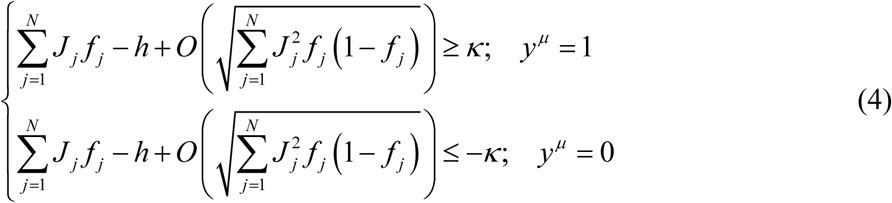

To guarantee *O*(1) capacity in the large *N* limit, it is necessary for the deviation of the average input from the threshold, 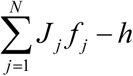, and the robustness parameter, *κ*, to be of the same order as (or less than) the width of the input distribution, 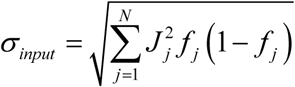. If not, input to the neuron will rarely cross the threshold (if the first conditions is violated) or the robustness margins (if the second condition is violated), and capacity for robust associative memory storage will be close to zero. Therefore,

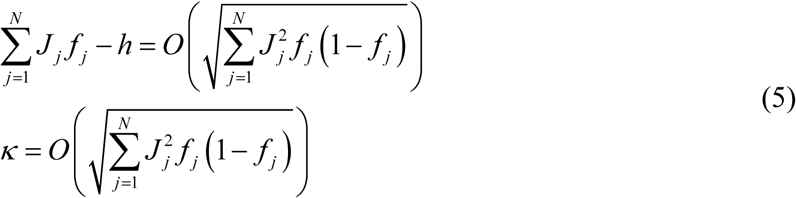

For two classes of neurons, one excitatory and one inhibitory, with similar weight magnitudes, Eqs. (5) give rise to various plausible scenarios for scaling of connection weights and robustness parameter with the network size:

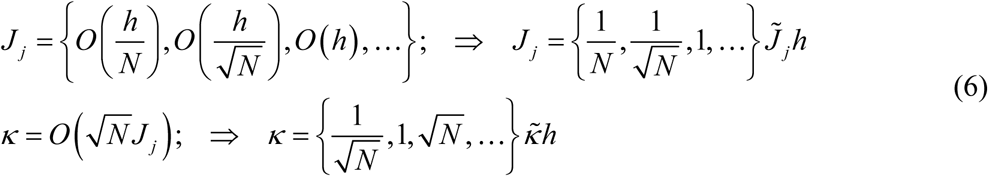

The normalized weights, 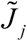, and the normalized robustness parameter, 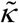, in Eqs. (6) do not scale with *N*.

The first scenario is usually used in associative memory models in conjunction with the replica theory (see e.g. ^5^):

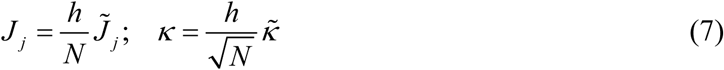

The second scaling scenario is traditionally used in balanced network models (see e.g. ^9^):

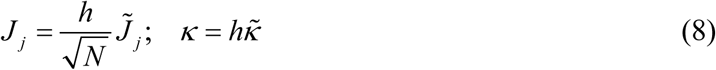

The third and the subsequent scenarios, in which *J* does not scale with *N*, or *J* increases with *N*, can be ruled out because they are biologically unrealistic. What is more, one can see from Eqs. (2) that the firing threshold in these cases can be disregarded, and the results of replica calculation become identical to the second scaling scenario.

In all models, scaling of *w^v^* is assumed to be the same as that of J, i.e. 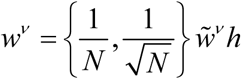. Substituting the normalized variables into Eqs. (2) we arrive at two problems, both governed by the same set of intensive parameters, *f_j_*, *f_out_*, 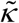, 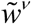, 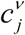, *g_j_*, as well as parameters *m, N*, and *k*:

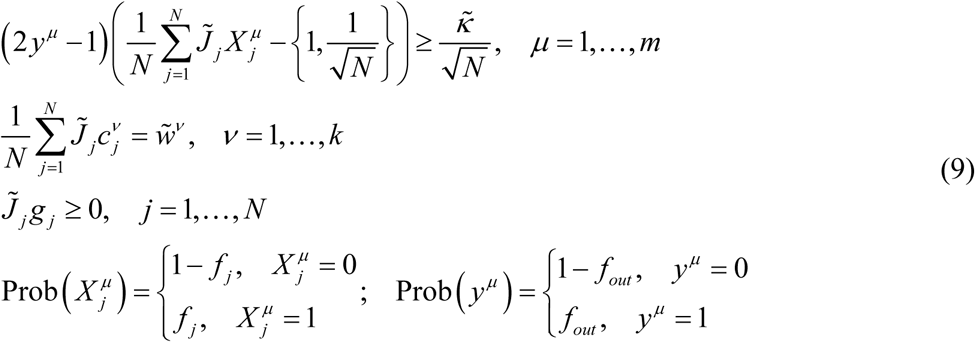

Note that the two formulations only differ in the threshold term.

### Replica theory solution of the model in the large *N* limit

We solve the above two models concurrently by following the steps of the replica theory solution outlined in ^7,8^. We begin by calculating the volume of the connection weight space, 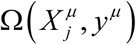, in which Eqs. (9) hold for a given set of associations:

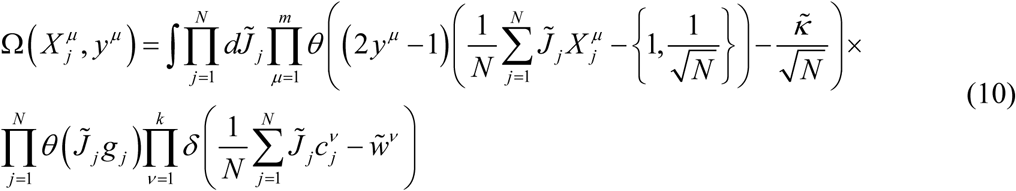

The typical volume of the solution space, *Ω_typical_*, is defined through the average of 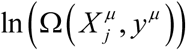 over the set of associations and is calculated by introducing *n* replica systems,

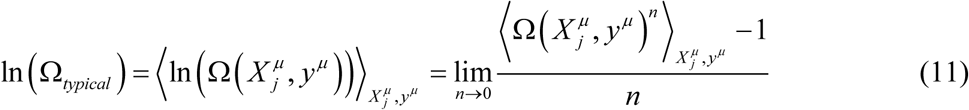

The quantity 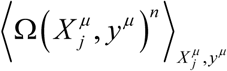 is then expressed as a single multidimensional integral:

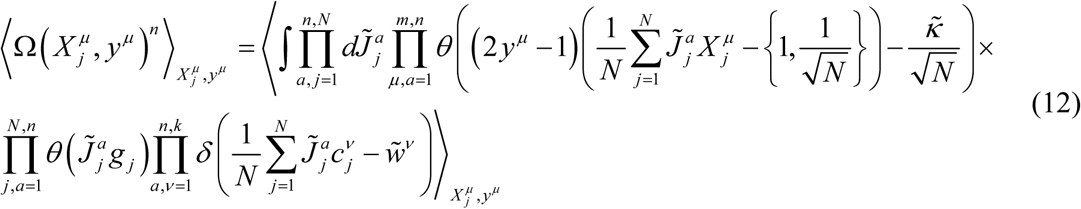

Input and output associations, 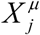 and *y^μ^*, are decoupled through the introduction of a new variable, 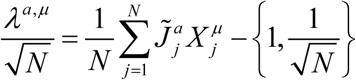:

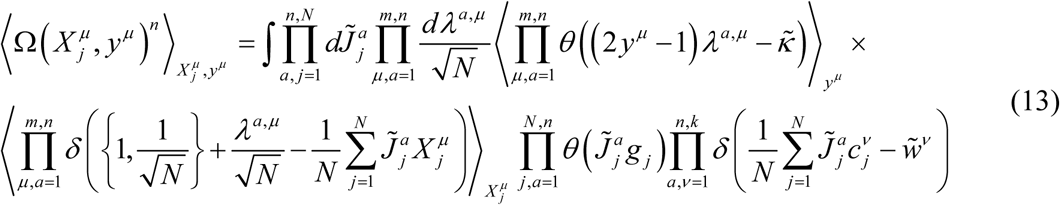

Next, the step-functions and the *δ*-functions are replaced with their Fourier representations, i.e.:

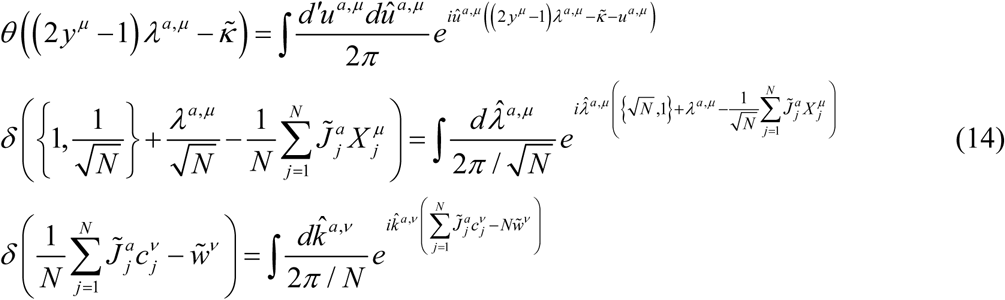

Symbol *d’* in these expressions and thereafter is designated for 0 to ∞ integration, whereas *d* is used to denote integration from −∞ to ∞.

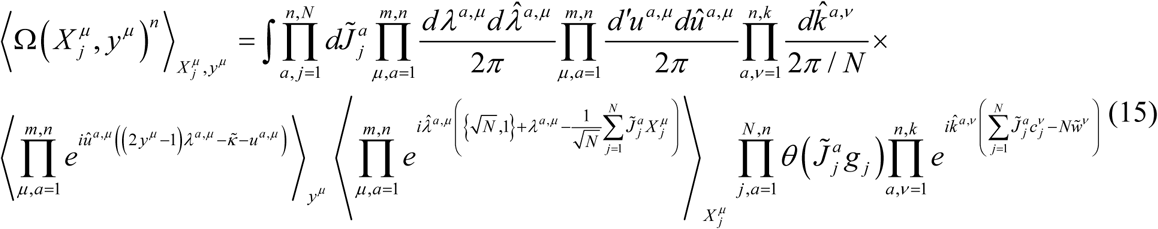

After averaging over the associations we arrive at:

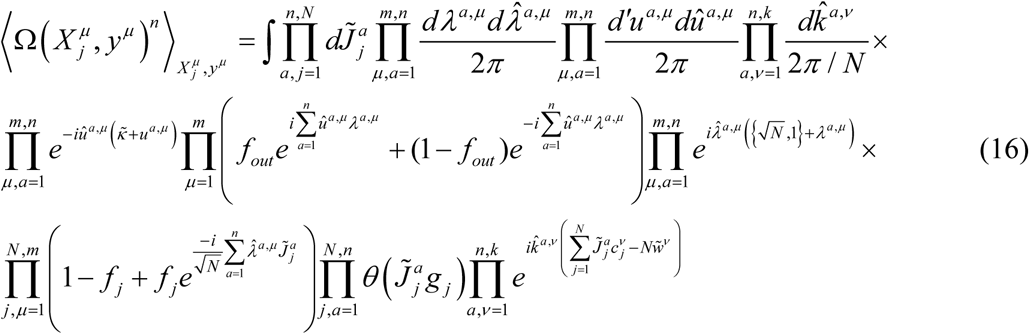

Replacing the argument of the first product in line three of Eq. (16) with an exponential expression that approximates it up to the second order in 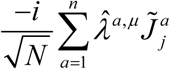, we obtain:

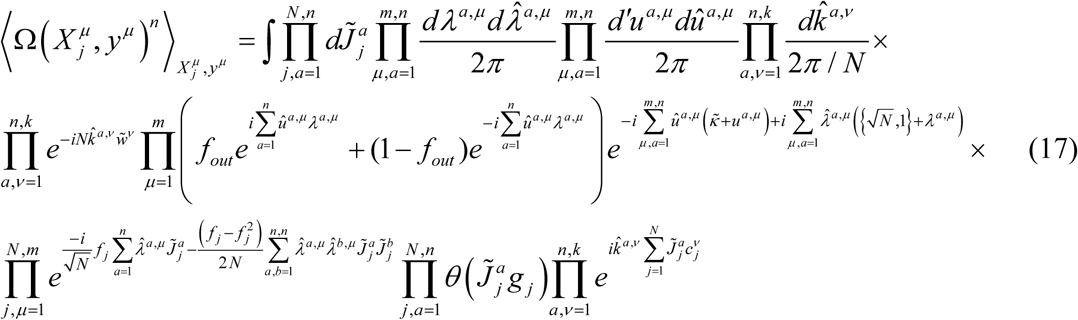

At this point, we introduce two sets of order parameters which allow us to decouple the products containing indices *j* and *μ*,

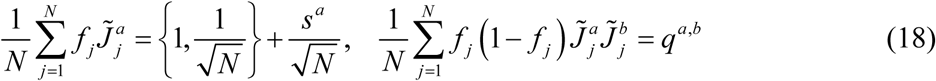

Insertion of these order parameters into Eq. (17) leads to the following expression:

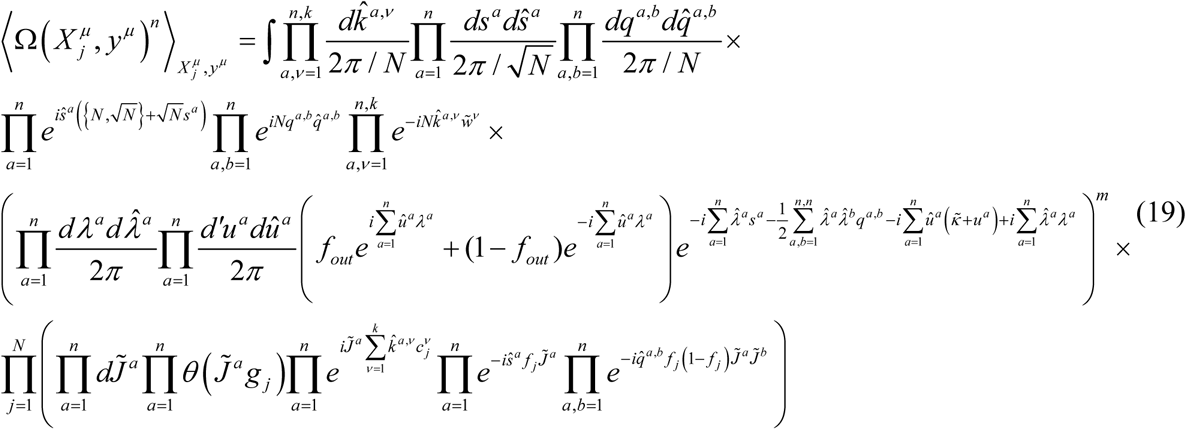

After the integration over 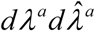 we arrive at:

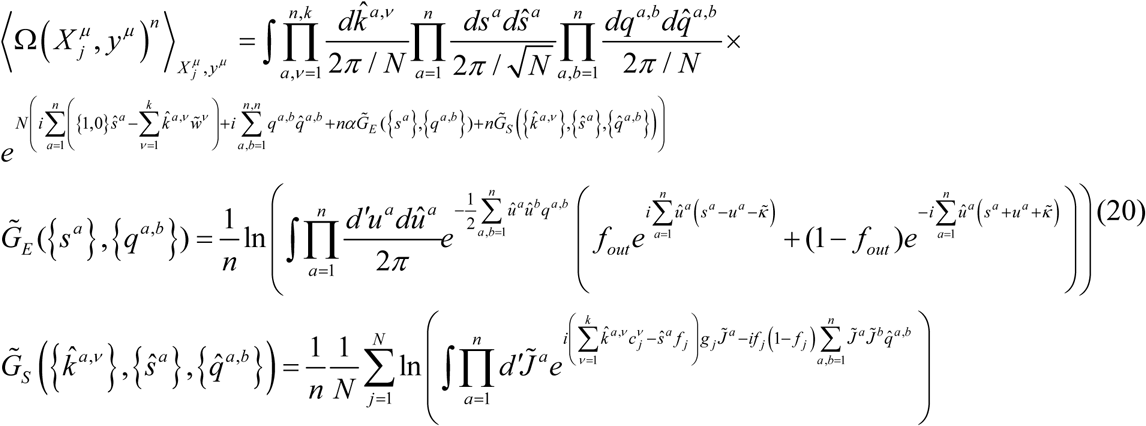

The integral in the first line of Eqs. (20) is calculated by using the steepest descent method combined with the assumption of a replica-symmetric saddle point, *s^a^ = s*, *q^a,a^ = q*_0_*, q*^a≠h^ *= q*, 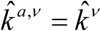, 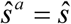, 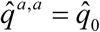, and 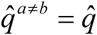:

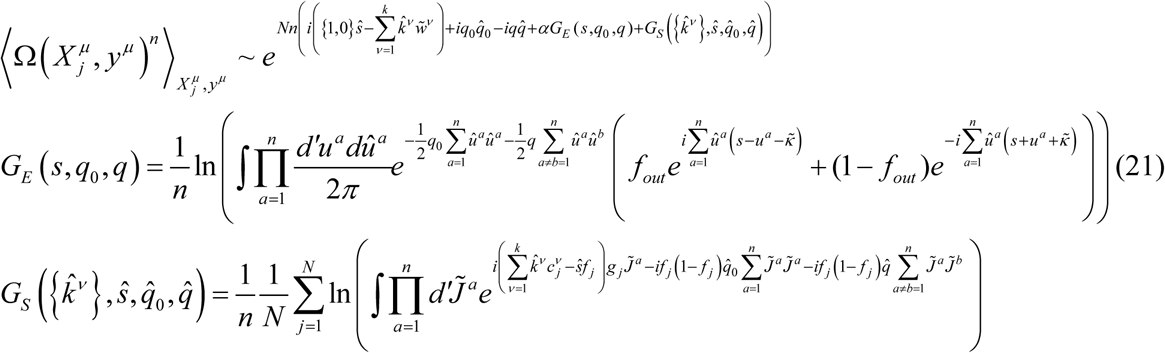

The non-redundant, replica-symmetric saddle point coordinates 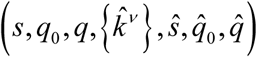 satisfy the following system of equations:

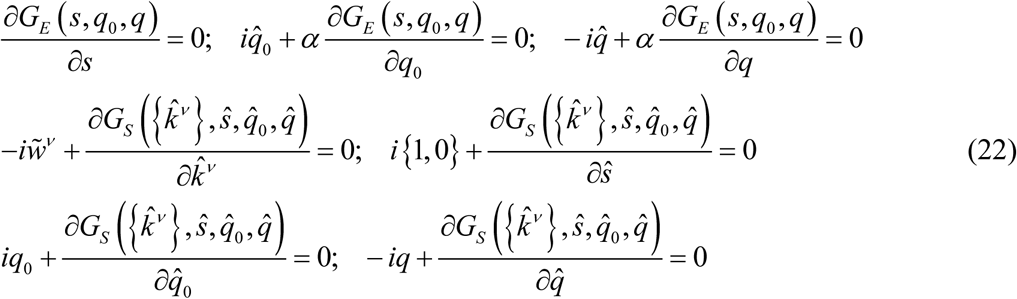

To simplify the expressions for *G_E_* and *G_s_*, we employ the Hubbard-Stratonovich transformation in the form of,

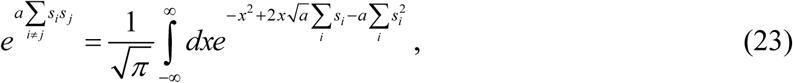

and take the *n* → 0 limit:

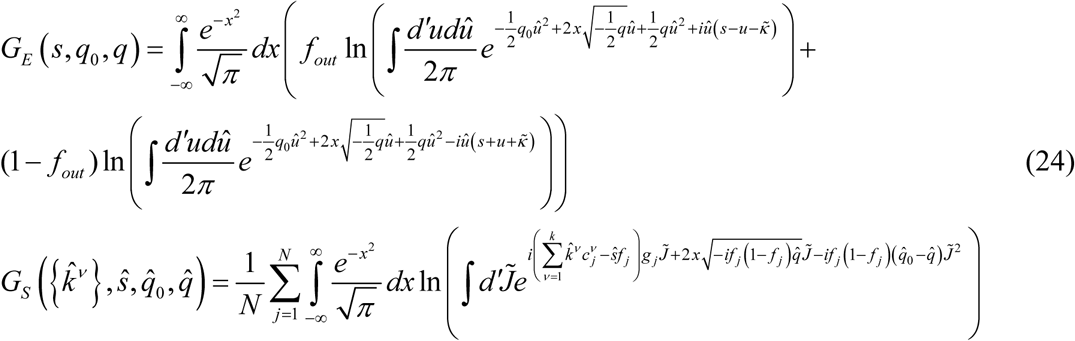

After calculating the integrals inside the arguments of the natural logarithms we obtain:

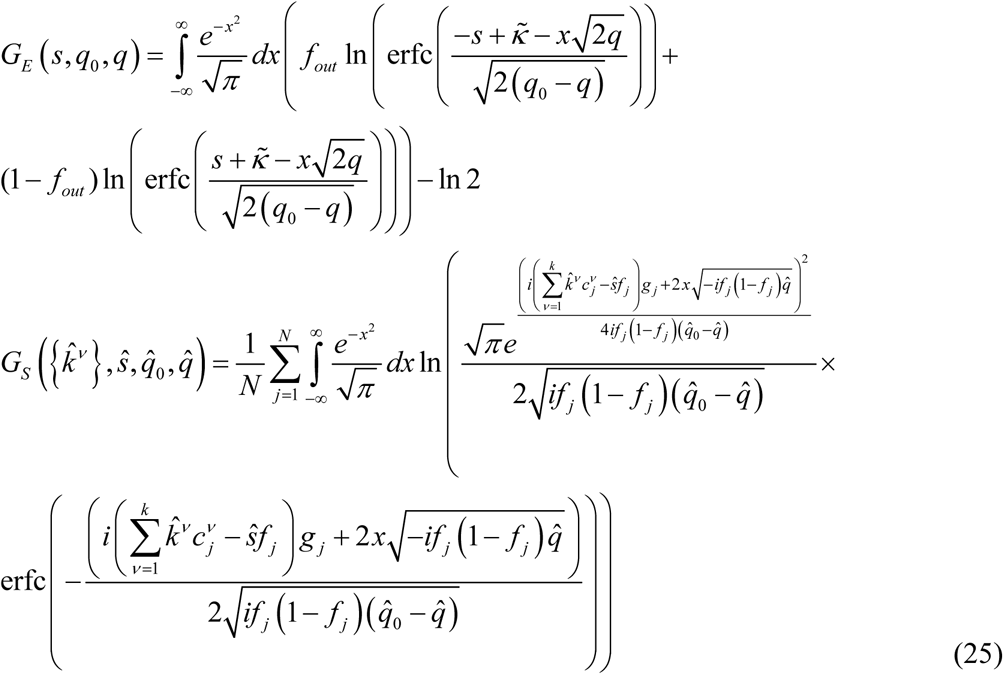

Next, we make substitutions, 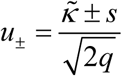, 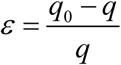, 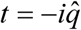, 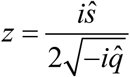, 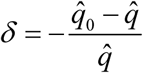, 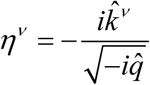, and rewrite *G_E_, G_s_*, and the saddle-point equations in terms of the new variables:

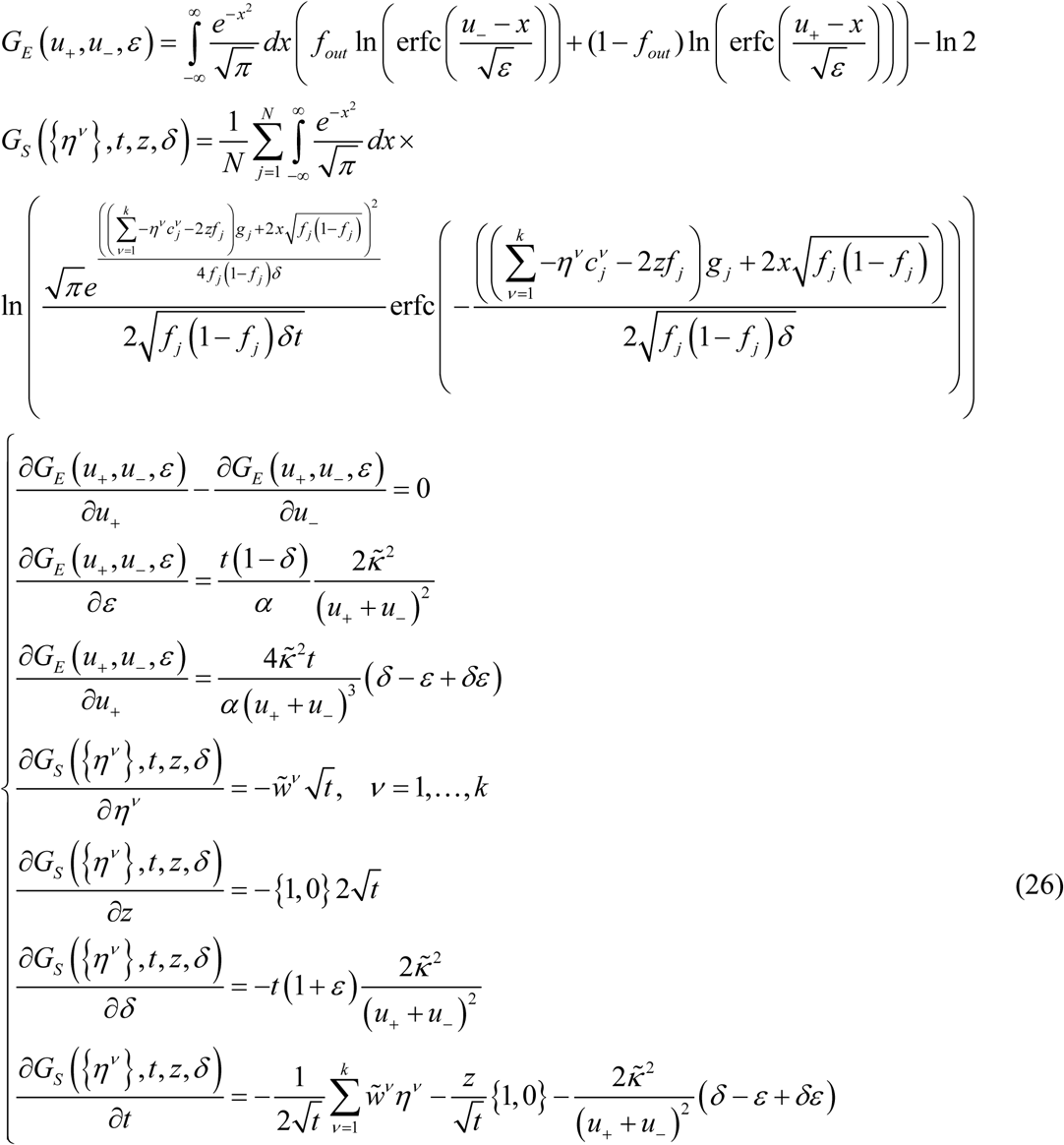

### Replica theory solution at critical capacity

At critical associative memory storage capacity, the saddle point Eqs. (26) can be simplified because as Ω*_typical_* tends to zero, *q*_0_ *− q* goes to zero as well. In this limit, parameters *ε* and *δ* are small, and the *k* + 6 saddle point equations can be expanded asymptotically to the leading orders in 1/ *ε* and *1/δ:*

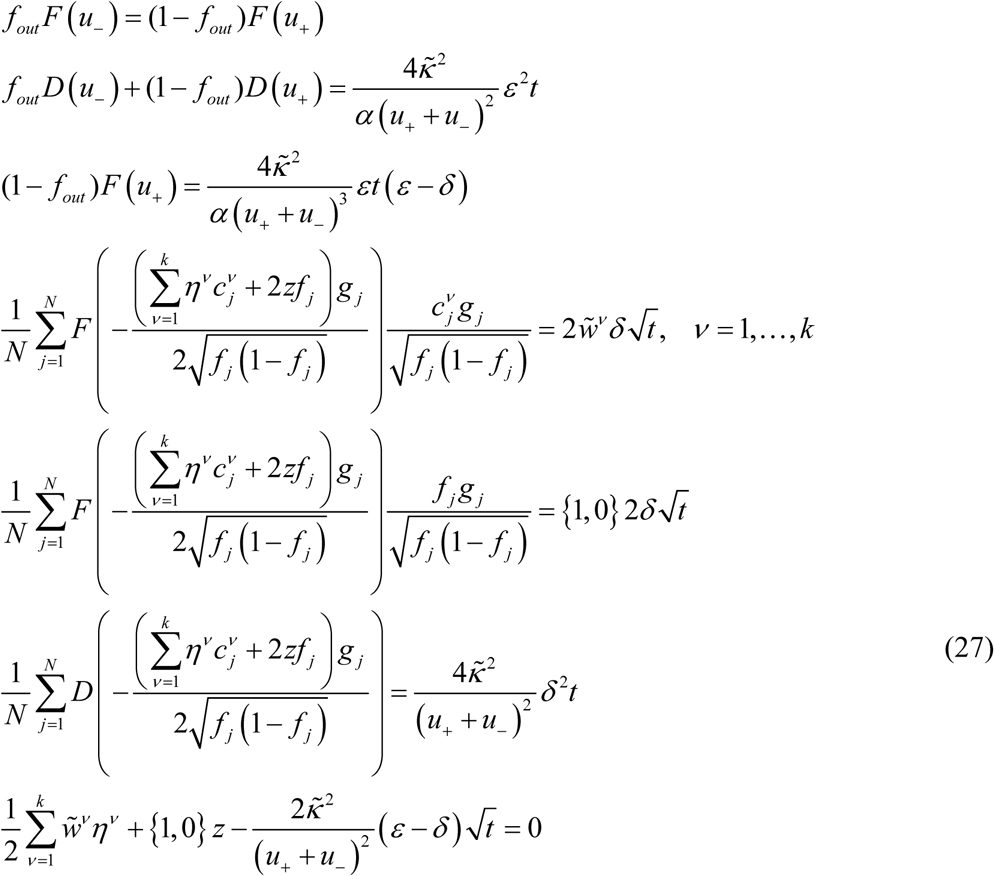

Special functions *E, F*, and *D* are introduced in the above expressions for conciseness:

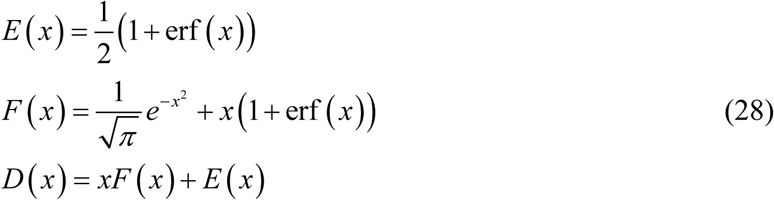

After eliminating *t, δ*, and *ε* from Eqs. (27) we arrive at the final result:

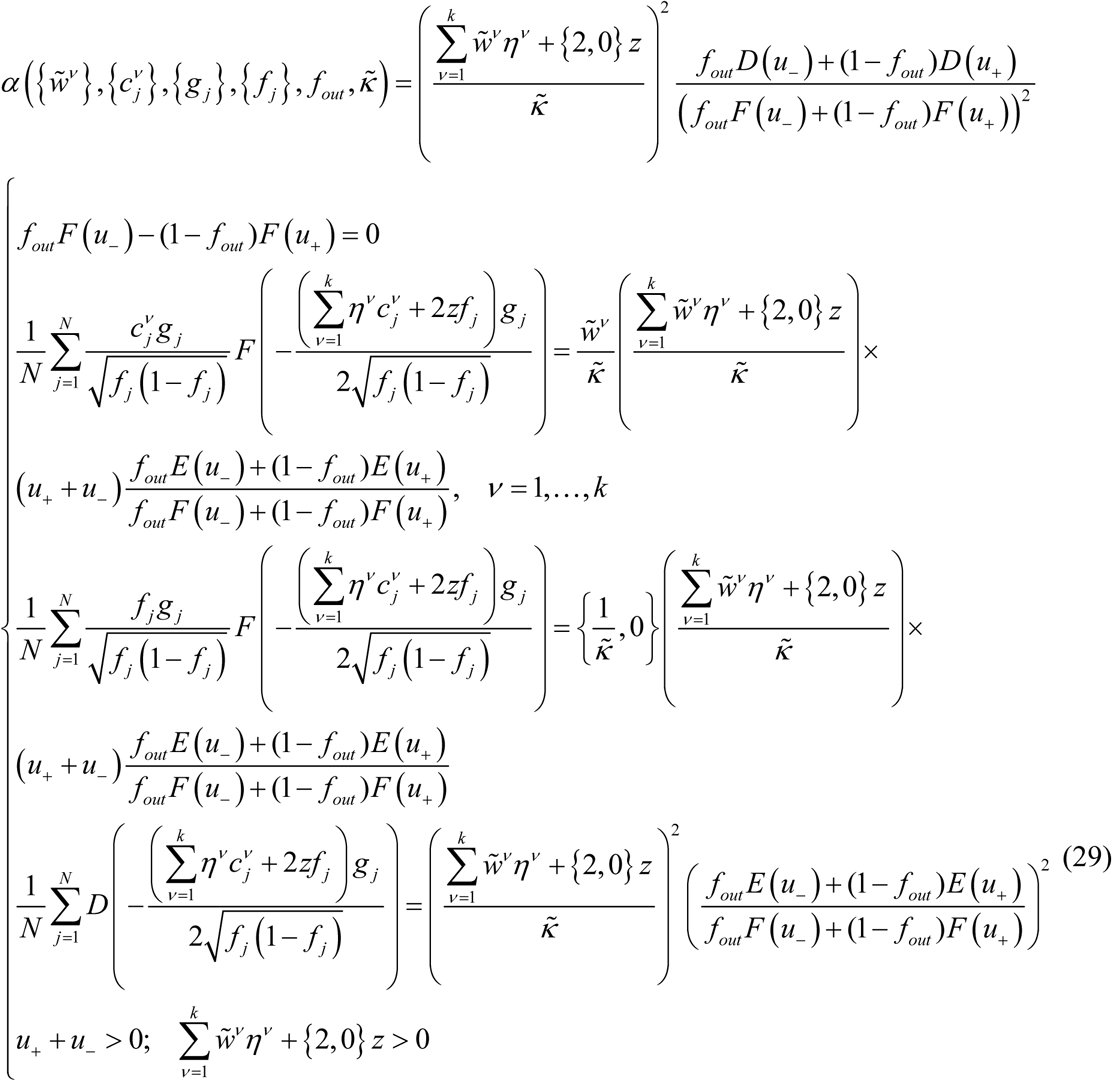

We note that to have a non-zero capacity, the number of input classes must be greater than the number of homeostatic constraints, *k*.

### Distribution of input weights at critical capacity

The probability density of input weights can be derived from the following general expression:

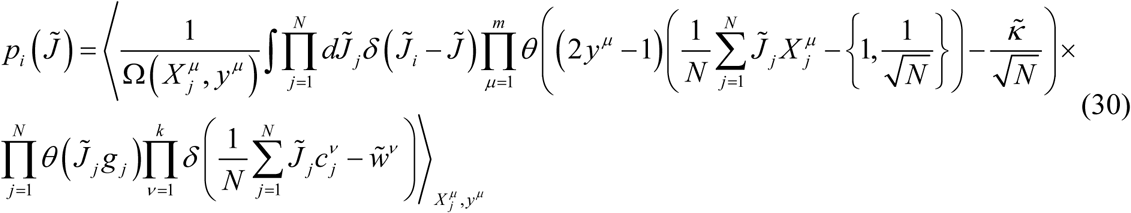

Eq. (30) can be cast in a form that closely resembles Eq. (12), allowing us to exploit the results of previous section. To that end, we introduce *n* replicas and take the limit of *n →* 0 after averaging over the associations:

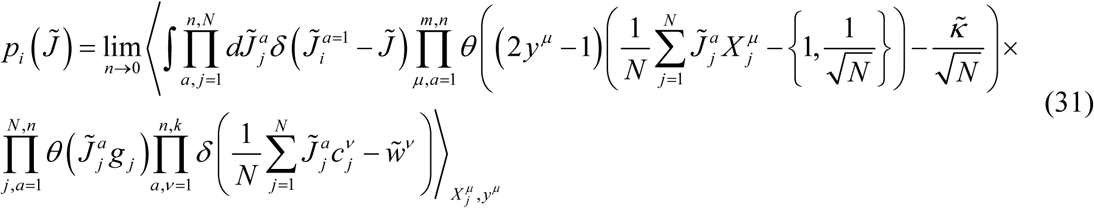

Following the steps outlined in Eqs. (12–22) we arrive at:

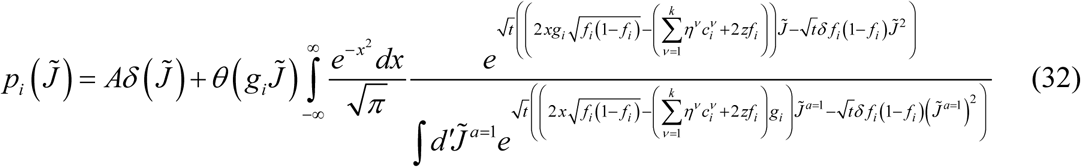

Constant *A* in this expression is defined by the normalization condition.

At critical capacity, the integrals in Eq. (32) can be calculated with the Laplace’s method, resulting in the following expression:

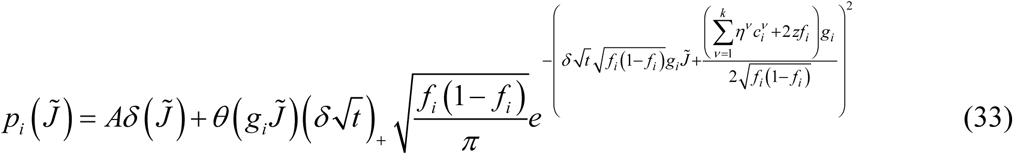

After substituting 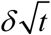 from the replica equations, Eqs. (27), and determining *A* from the normalization condition, we obtain the distributions of connection weights for different input classes:

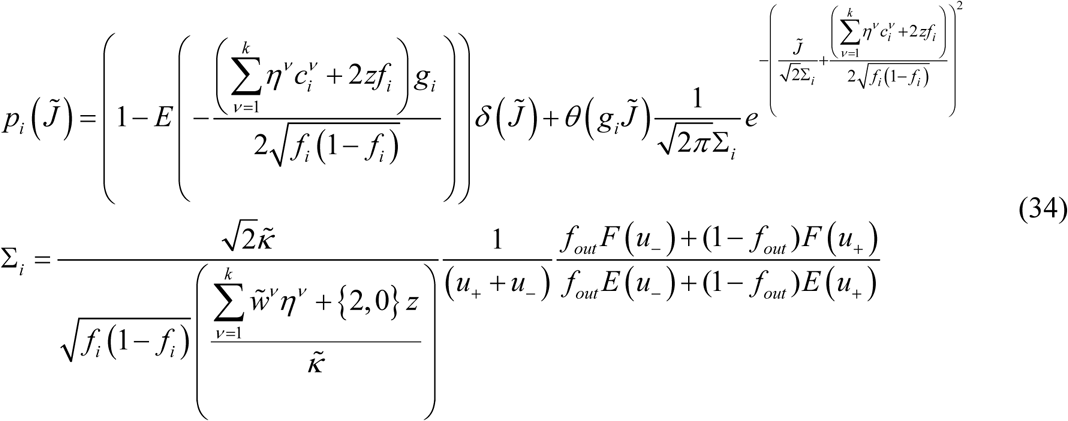

These distributions are composed of Gaussian functions truncated at zero and finite fractions of zero-weight connections. Parameters Σ*_i_* describe the distribution widths.

Connection probabilities and distributions of non-zero connection weights follow from Eqs. (34):

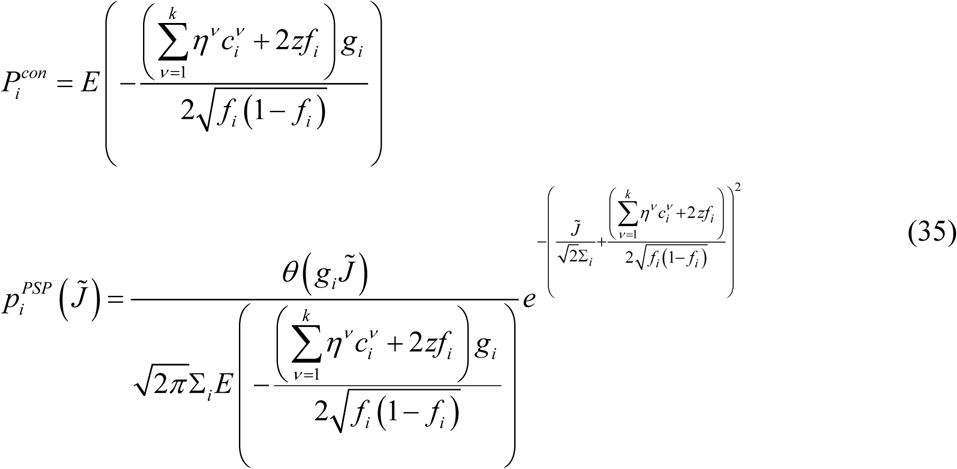

### Solution in the high-weight regime considered in the main text

In the main text, we consider a simplified problem in which there are only two classes of neurons, one inhibitory and one excitatory, each neuron has a single *l*_1_-norm constraint on the weights of its inputs (*v =* 1, 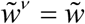, 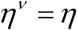, 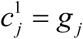), and the firing probabilities are homogeneous (*f_j_ = f_out_ = f*). In this case, we introduce new variables, 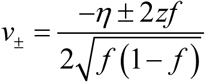 and 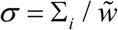 and the general solution of Eqs. (29, 35) simplifies significantly:

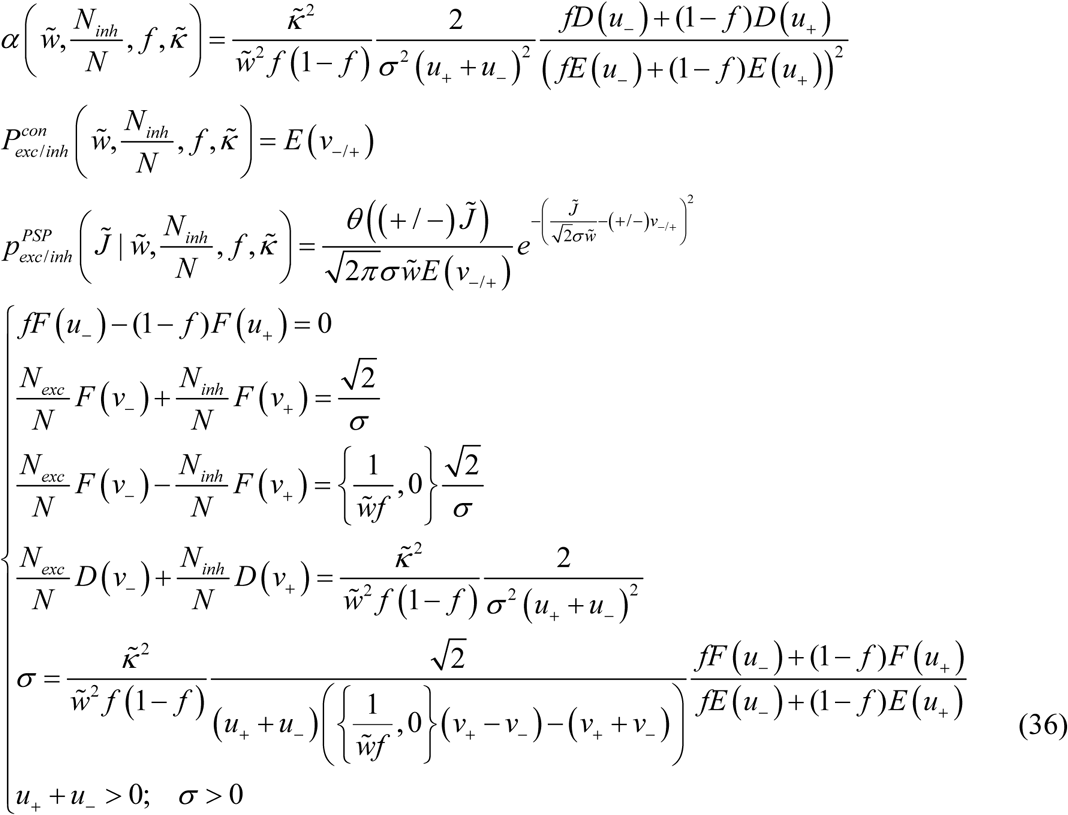

Note that the difference between the replica solutions of the associative and balanced models explicitly appears only in two places (curly brackets) in Eqs. (36).

These equations make it clear that the solution of the associative model in the high-weight limit, 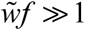, converges to the solution of the balanced model. However, since the value of 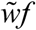 estimated from experimental data is large but finite (see the main text), we also examined the agreement between the results of the two models by solving Eqs. (36) numerically for different values of 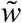 and 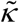. Figure S1 shows that for values of 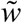 in the [10 100] range, results of the two models agree within 10%, and the agreement improves with increasing 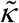. What is more, in the limit of high-weight the solution depends only on 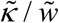 (straight isocontour lines in Figure S1), or alternatively, on a parameter 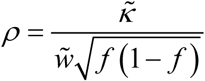. The latter was introduced by Brunel, et al ^5,6^ and is referred to as the rescaled robustness. Parameter *ρ* can serve as a proxy for the ratio of robustness and standard deviation in postsynaptic input, *κ*/ *σ_input_*. With this, Eqs. (36) simplify further:

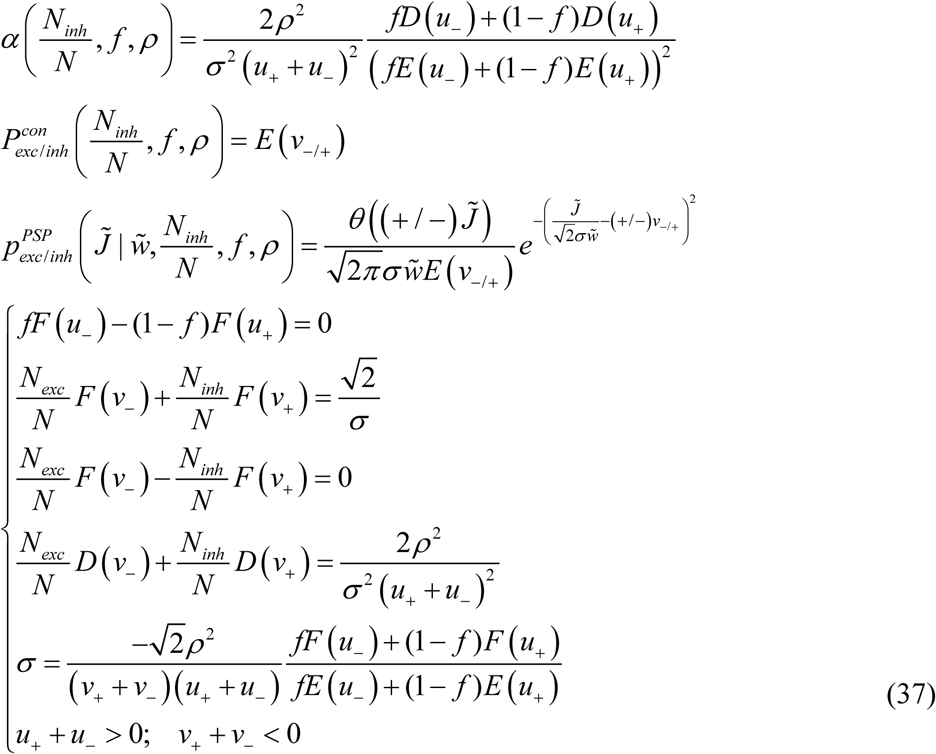

We note that since 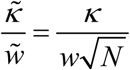 for both models, rescaled robustness, *ρ*, and Eqs. (37) are model independent.

The fact that the solutions of the two models converge in the high-weight regime is not surprising. In this regime, *Nwf* ≫ *h*, and, as a result, mean excitatory and inhibitory inputs to the neuron are much greater than the threshold of firing. One can show that in this case *h* in Eqs. (2) can be disregarded, and the solution becomes independent of scaling of *J* with *N*.

### Numerical solution for finite *N*

The problem of Eqs. (2) is convex, and hence, it can be solved numerically within the standard constrained optimization framework. Numerical solutions are obtained for finite networks, and the results are independent on the assumptions of scaling of model parameters with *N*.

Below, we consider two learning scenarios: (*i*) feasible load, in which associations can be learned with specified robustness, and (*ii*) non-feasible load, in which the number of presented associations is so large that Eqs. (2) have no solution.

In the case of feasible load, the region of solutions is nonempty, and one must employ additional considerations to limit the results to a single, “optimal” solution. We do this by choosing the solution that minimizes 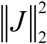,

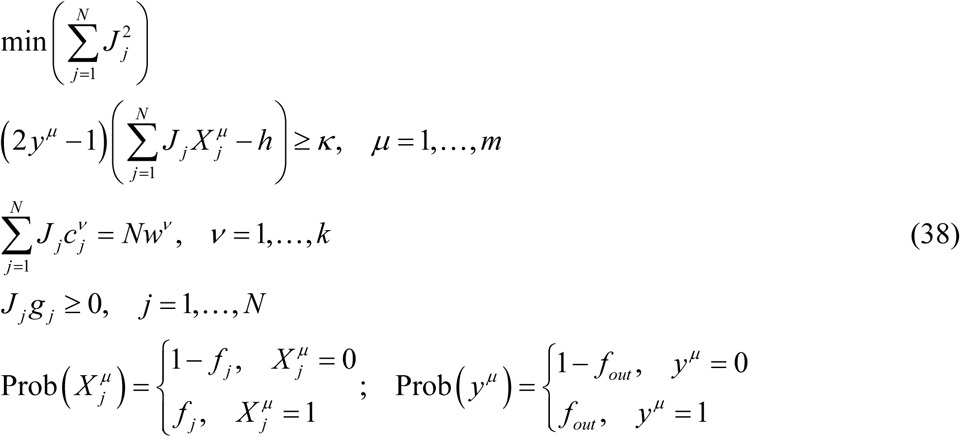

In the non-feasible case, we choose the solution that minimizes the sum of deviations, *s^μ^*, of not robustly learned associations from the corresponding margin boundaries. This solution can be obtained by solving the following linear optimization problem:

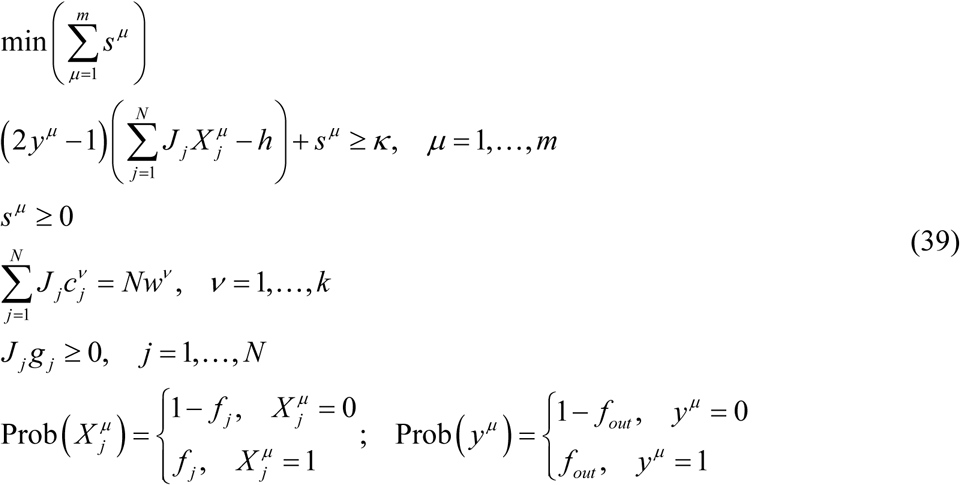

The problems outlined in Eqs. (38, 39) were solved in MATLAB in the following sequence of steps. Given the associative memory load, *α = m* / *N*, we first solved the problem of Eqs. (39), utilizing the *linprog.m* function, to find the distances, *s^μ^*. If some of these distances are greater than zero, the problem is non-feasible. If all *s^μ^ =* 0, the problem is feasible, in which case we used connection weights resulting from Eqs. (39) as a starting configuration and solved Eqs. (38) by using the *quadprog.m* function.

Figure S2 shows the results of numerical simulations for neurons loaded to capacity at different values of rescaled robustness. Results for *N* = 200, 400, and 800 inputs are shown together with the replica theory solution (*N* → ∞). With increasing *N*, the numerical solutions gradually approach the results of the replica theory, which serves as an independent validation of numerical and theoretical calculations.

The dependence of structural and dynamical properties of associative networks of *N* = 200, 400, and 800 neurons on memory load and rescaled robustness is shown in Figures S3, S4, and S6. These maps do not depend strongly on *N*, suggesting that the network properties would not change significantly if the network size was increased further, e.g. to a few thousand neurons.

### Numerical values of model parameters and results

In this section, we provide values of various parameters related to network structure and dynamics. This is done for the two network settings corresponding to the red and green asterisks in Figures 2-5 of the main text.

**Table S1:** Input and output model parameters corresponding to the red and green asterisks in Figures 2-5 of the main text.

|  | Parameter name | Red asterisk | Green asterisk |
| --- | --- | --- | --- |
| Input parameters | Number of neurons, $N$ | 800 | 800 |
| | Inhibitory neuron fraction, $N_{inh}/N$ | 0.20 | 0.20 |
| | Firing probability, $f$ | 0.20 | 0.20 |
| | Threshold of firing, $h$ | 20 mV | 20 mV |
| | Scaled average absolute connection weight, $\tilde{w}$ | 70 | 70 |
| | Scaled memory load, $\alpha/\alpha_c$ | 0.90 | 0.90 |
| | Rescaled robustness, $\rho$ | 0.50 | 1.3 |
| Output parameters | Memory load, $\alpha$ | 0.38 | 0.20 |
| | Robustness parameter, $\kappa$ | 25 mV | 64 mV |
| | Excitatory connection probability, $P_{exc}^{con}$ | 0.26 | 0.14 |
| | Inhibitory connection probability, $P_{inh}^{con}$ | 0.66 | 0.46 |
|  | CV of excitatory connection weights | 0.89 | 0.99 |
|  | CV of inhibitory connection weights | 0.75 | 0.86 |
| | Average number of steps to limit cycle | 32 | $2.3 \times 10^4$ |
|  | CV of ISI | 0.67 | 0.88 |
|  | Spike cross-correlation coefficient | 0.24 | 0.09 |
| | Excitatory input (mean ± s.d.) | $133 \pm 14$ mV | $125 \pm 38$ mV |
| | Inhibitory input (mean ± s.d.) | $-144 \pm 35$ mV | $-155 \pm 48$ mV |
| | Total input (mean ± s.d.) | $-11 \pm 38$ mV | $-30 \pm 60$ mV |
|  | Exc.-inh. input correlation coefficient | -0.96 | -0.79 |
|  | Sequence retrieval probability | 0.08 | 1 |
| | Noise tolerance, $\sigma_{noise}/\sigma_{input}$ | 0 | 0.35 |

Figure S7 illustrates partial cancellation of excitatory and inhibitory inputs and the relationships among the total input, firing threshold, and robustness parameter based on the values from Table S1.

---

**Figure S1:**
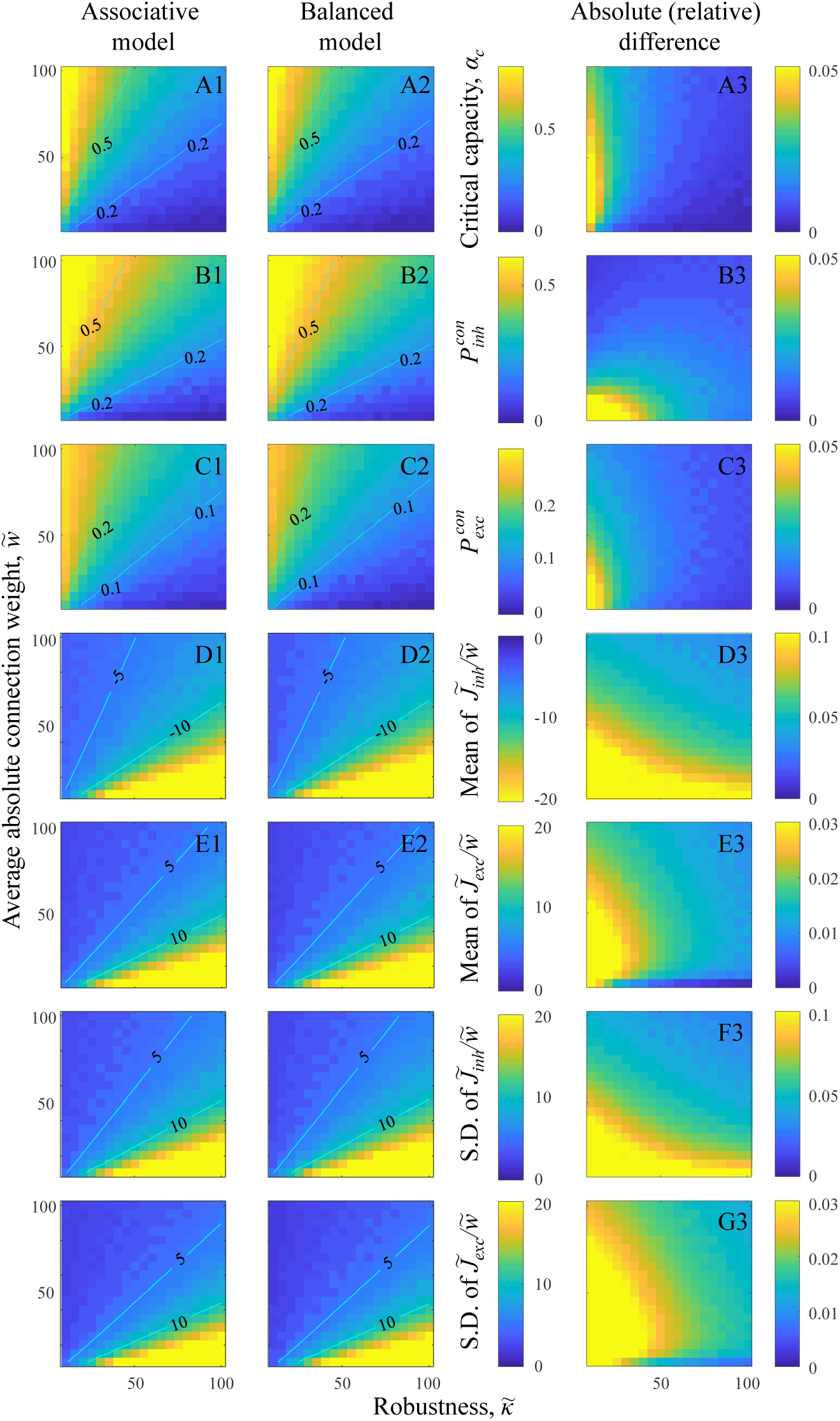
Theoretical solutions for the associative and balanced models converge in the limit of 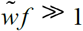. In this limit, model results depend only on 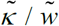, in agreement with Eqs. (37). **A.** Maps of critical capacity as functions of 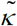 and 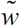 for the associative (A1) and balanced (A2) models. Straight isocontours confirm that the results depend only on 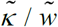. Absolute difference of the two maps (A3) shows that critical capacities of the two models converge in the limit of 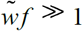. Same for the probabilities of inhibitory **(B)** and excitatory **(C)** connections. **D.** Maps of mean, non-zero, inhibitory connection weights as a function of 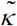 and 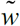 for the associative (D1) and balanced (D2) models, as well as the absolute relative difference of these maps (D3). Same for the mean, non-zero, excitatory connection weights **(E)**, and standard deviations of non-zero inhibitory **(F)** and excitatory **(G)** connection weights.

**Figure S2:**
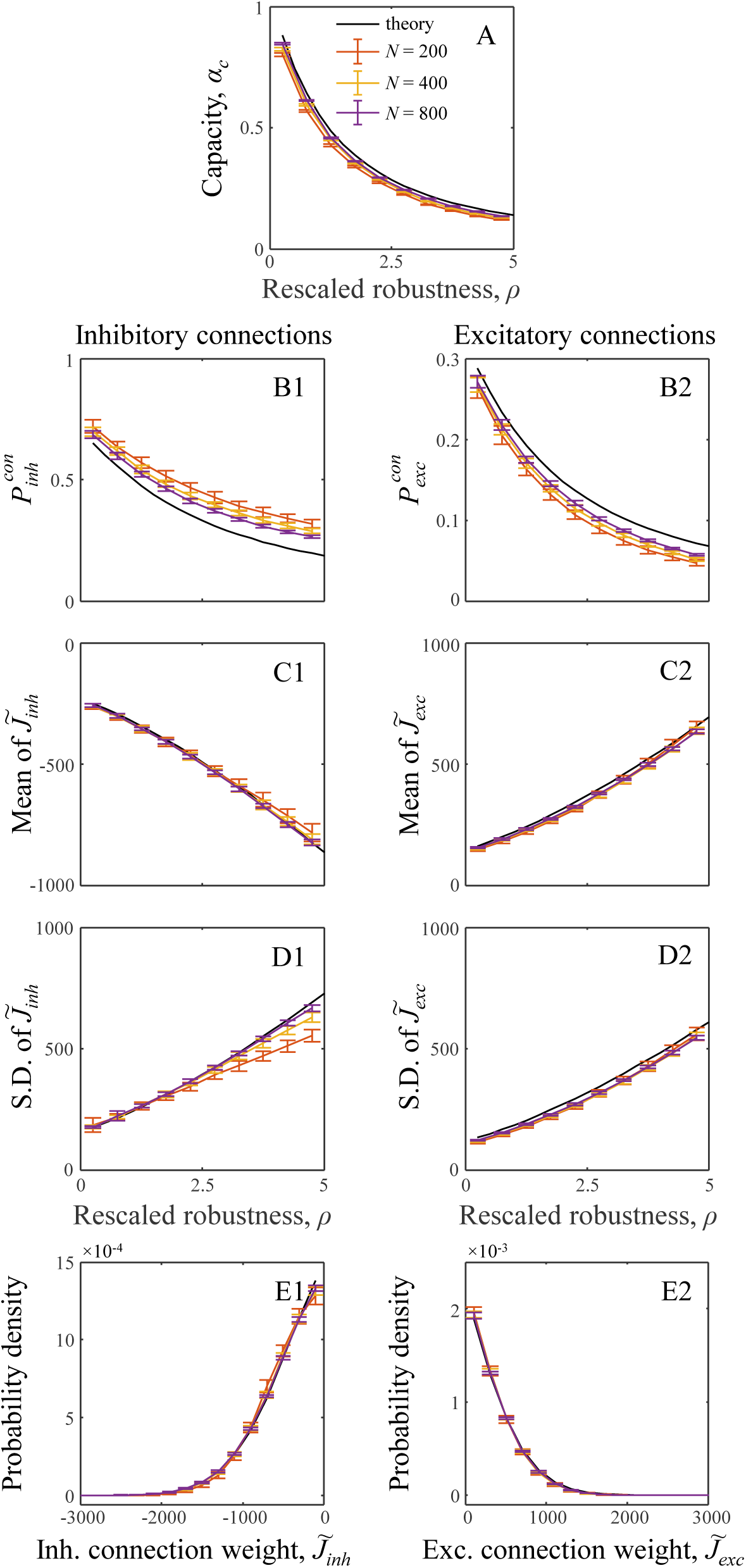
Validation of theoretical results of Eqs. (37) with numerical simulations performed for *N* = 200, 400, and 800 inputs. **A.** Capacity as a function of rescaled robustness, *ρ*. With increasing *N*, numerical results (error-bars) obtained with convex optimization [Eqs. (38, 39)] approach the theoretical solution (black line). Error-bars indicate standard deviations calculated based on 100*N* simulations. Same for the probabilities of non-zero inhibitory **(B1)** and excitatory **(B2)** connections, mean non-zero inhibitory **(C1)** and excitatory **(C2)** connection weights, and standard deviations of non-zero inhibitory **(D1)** and excitatory **(D2)** connection weights. Panels **(E1)** and **(E2)** illustrate the match between theoretical and numerical probability densities of non-zero inhibitory and excitatory connection weights for *ρ* = 1.

**Figure S3:**
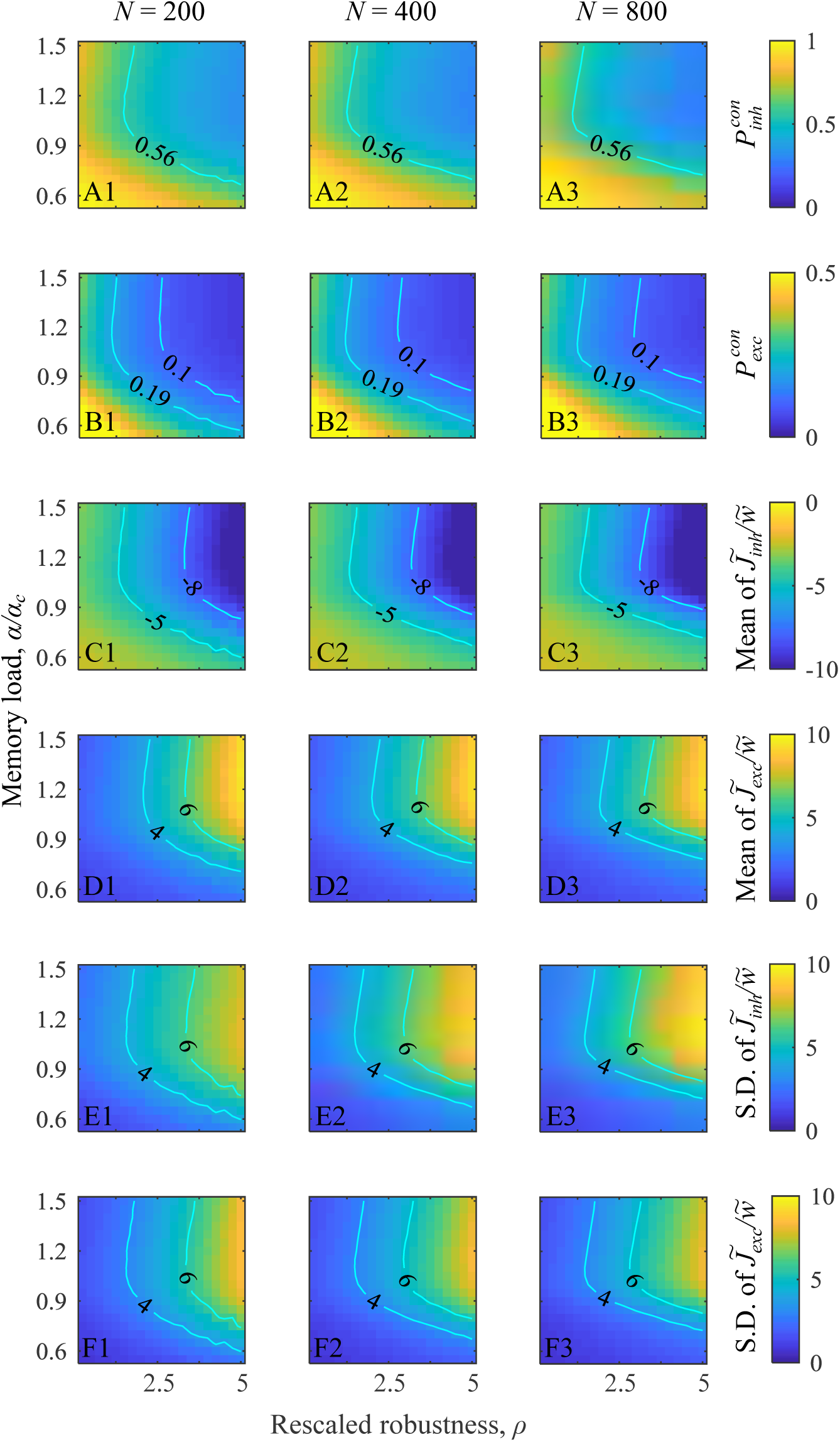
Properties of neuron-to-neuron connectivity in associative networks. **A.** Inhibitory connection probabilities as functions of rescaled robustness and relative memory load in networks of *N* = 200 (A1), *N* = 400 (A2), and *N =* 800 (A3) neurons. Same for excitatory connection probabilities **(B)**, means of non-zero inhibitory **(C)** and excitatory (**D**) weights (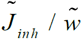 and 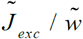), and standard deviations of inhibitory (**E**) and excitatory (**F**) weights. Isocontour lines match those used in the main text.

**Figure S4:**
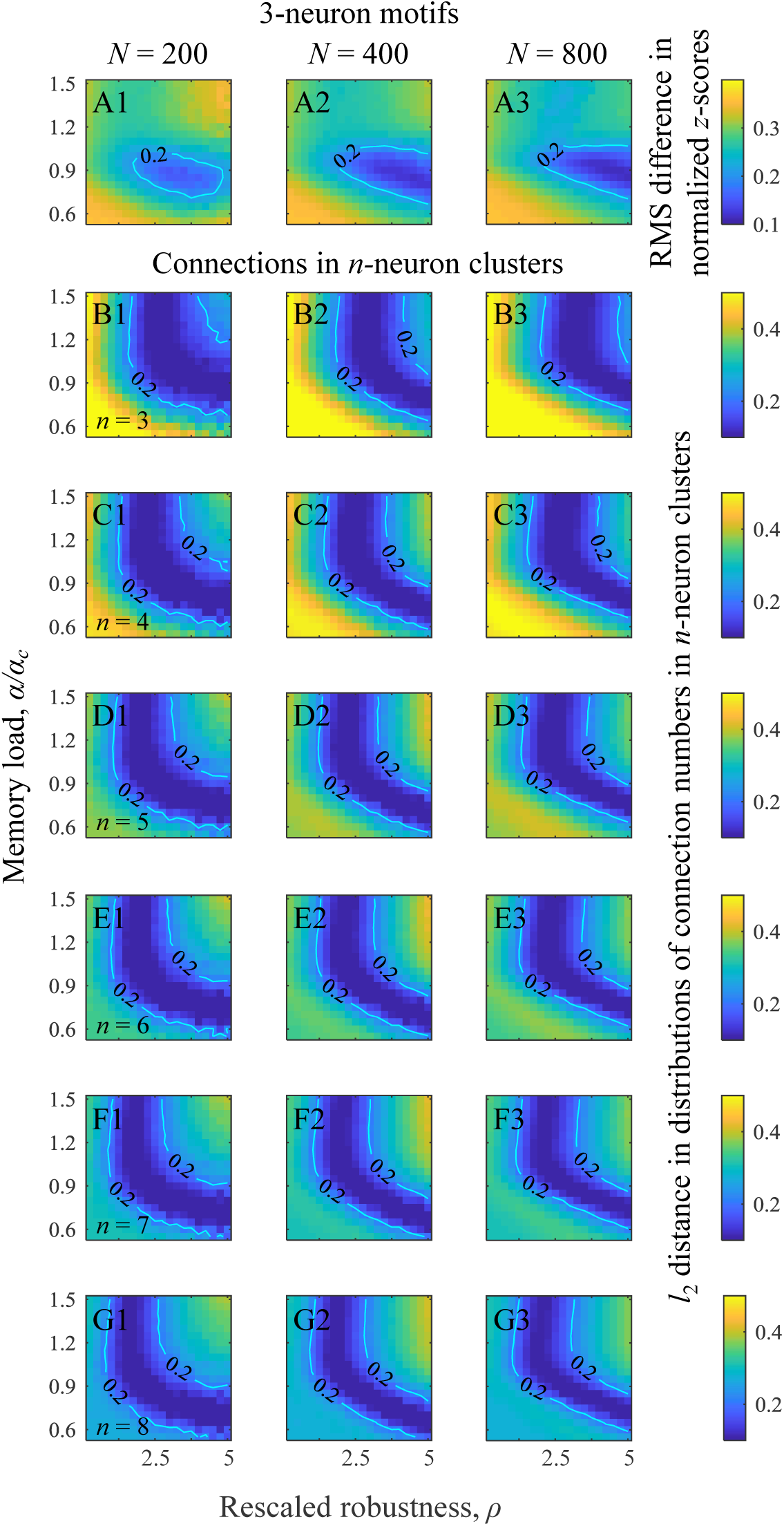
Dependence of higher-order structural properties of associative networks on network size. **A.** Maps of root-mean-square (RMS) differences between normalized *z-*scores of excitatory 3-neuron motifs observed in associative networks of *N* = 200, 400, and 800 neurons and those reported by the Blue Brain project ^13^. **B.-G**. Maps of *l*_2_ distances between distributions of connection numbers in 3–8 excitatory neuron clusters in associative networks of *N* = 200, 400, and 800 neurons and those observed experimentally ^14^. Isocontour lines match those used in the main text.

**Figure S5:**
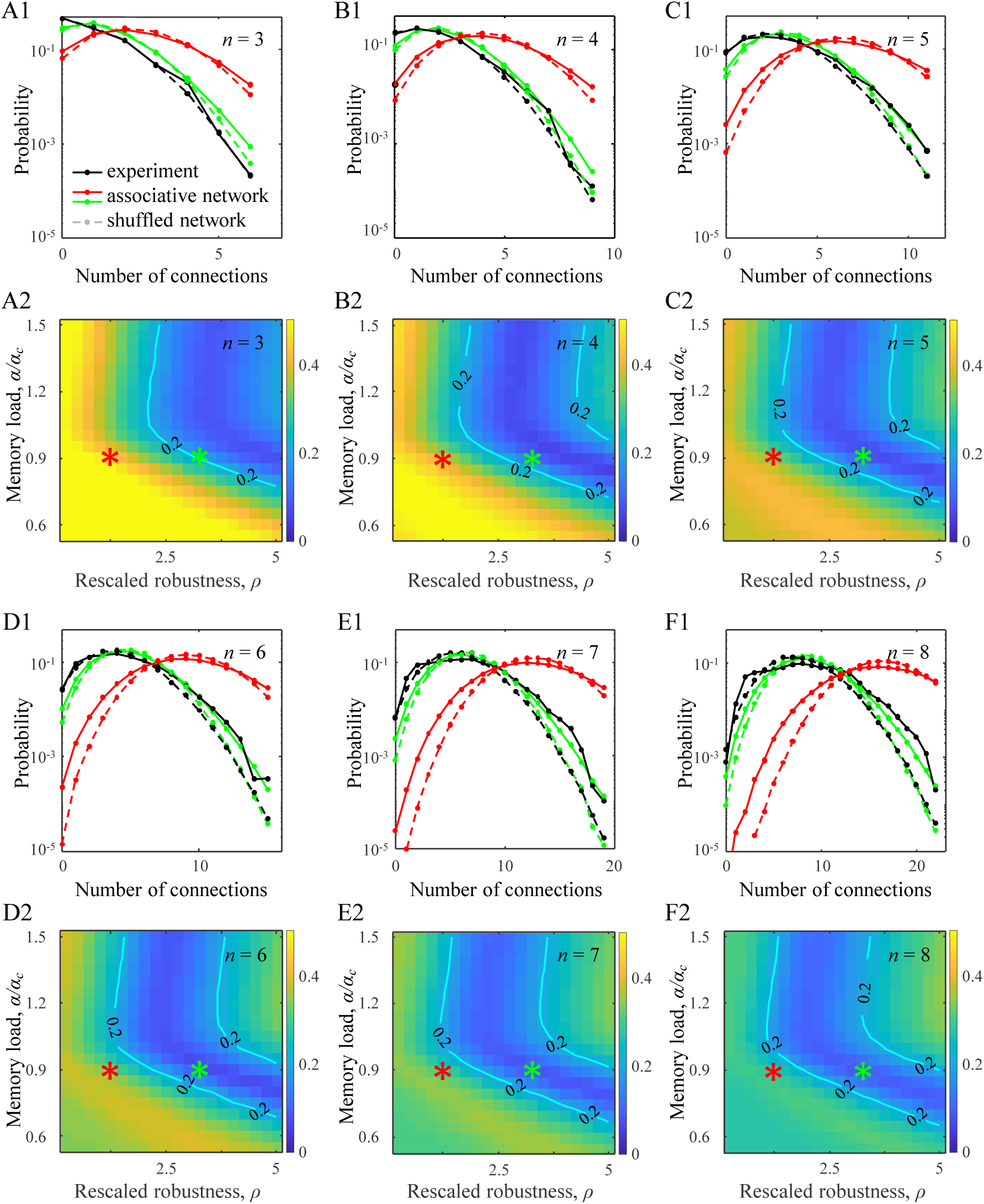
Distributions of non-zero connection numbers in clusters of 3–8 excitatory neurons in associative networks. **A1-F1**. Solid red and green lines illustrate distributions obtained in associative networks for the parameter settings indicated by the red and green asterisks. Solid black curves indicate the corresponding results for local cortical networks based on electrophysiological measurements ^2^. Dashed lines show distributions in randomly shuffled networks (see Methods for details). **A2-F2**. Maps of *l*_2_ distances between connection number distributions in associative and cortical networks ^2^. Numerical results of the associative model were generated based on networks of *N* = 800 neurons.

**Figure S6:**
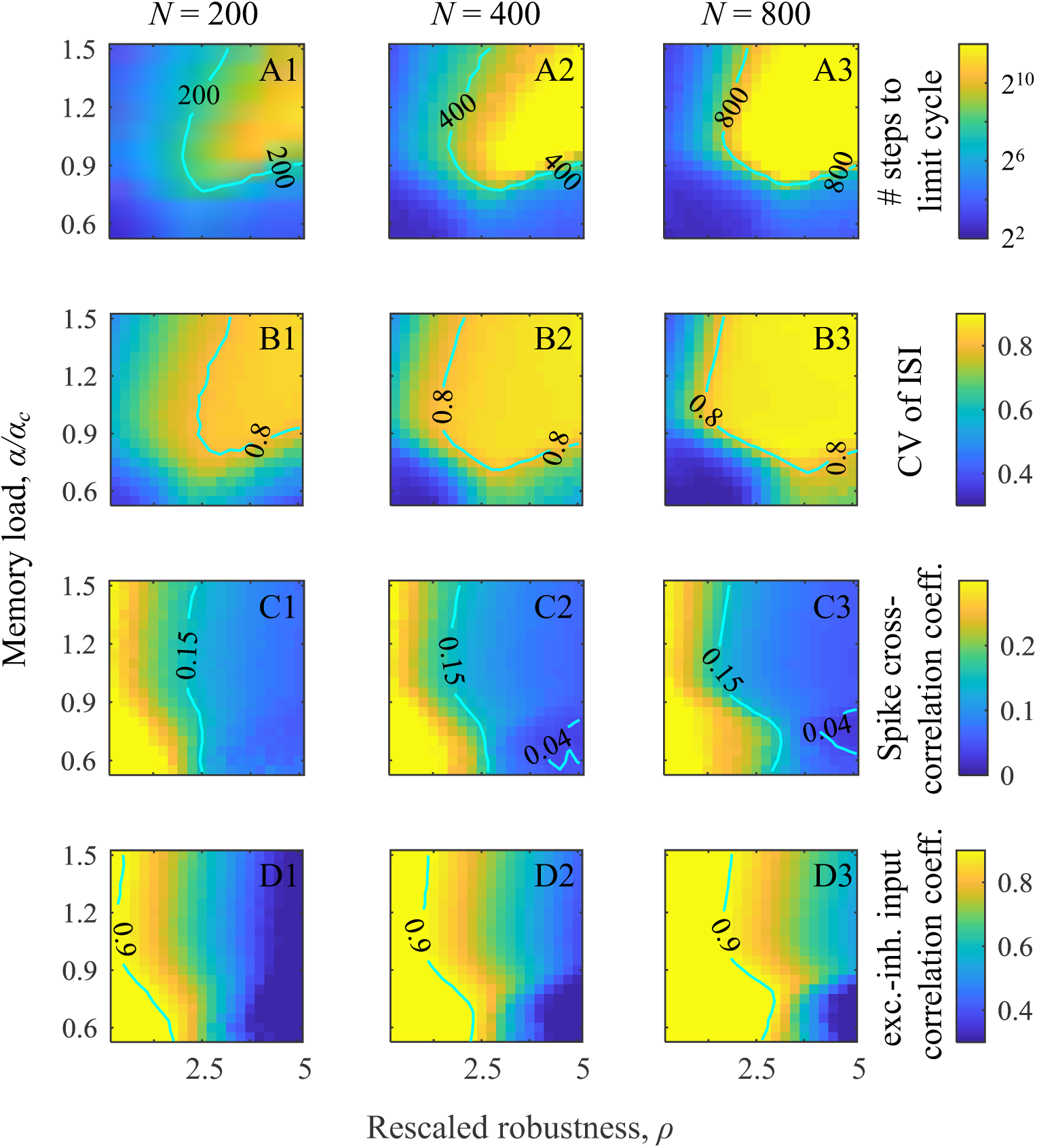
Dependence of dynamical properties of associative networks on network size. **A.** Maps for the average durations of transient network dynamics as functions of rescaled robustness and relative memory load for networks of *N* = 200, 400, and 800 neurons. Same for CV values of inter-spike-intervals (ISI) **(B)**, spike cross-correlation coefficients (**C**), and correlation coefficients of excitatory and inhibitory inputs (**D**). Isocontour lines match those used in the main text.

**Figure S7:**
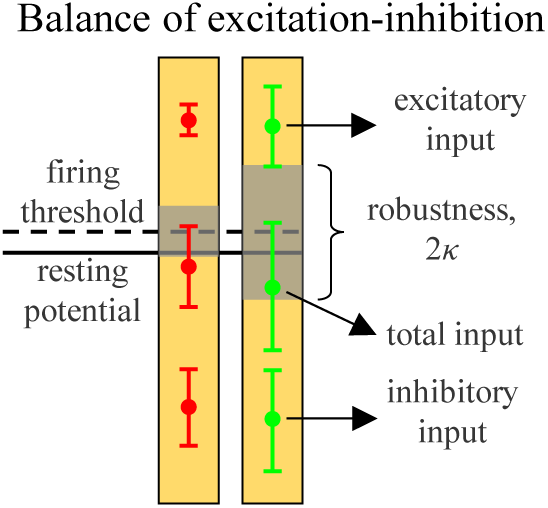
Excitatory and inhibitory inputs in relation to firing threshold and robustness. Left and right halves of the figure are based on the data from Table S1, and correspond to the red and green asterisks from Figures 2–5 of the main text. The average excitatory and inhibitory inputs are much larger than the threshold of firing. However, the total input lies within one standard deviation from the firing threshold due to a partial cancellation of its excitatory and inhibitory components.

